# Vicarious reward unblocks associative learning about novel cues in male rats

**DOI:** 10.1101/612697

**Authors:** Sander van Gurp, Jochen Hoog, Tobias Kalenscher, Marijn van Wingerden

**Affiliations:** Social Rodent Lab, Department of Experimental Psychology, Heinrich-Heine-University, Düsseldorf, Germany

## Abstract

Many species, including humans, are sensitive to social signals and their valuation is important in social learning. When social cues indicate that another is experiencing reward, they could convey vicarious reward value and prompt social learning. Here, we introduce a task that investigates if vicarious reward delivery in male rats can drive reinforcement learning in a formal associative learning paradigm. Using the blocking/unblocking paradigm, we found that when actor rats have fully learned a stimulus-self reward association, adding a cue that predicted additional partner reward unblocked associative learning about this cue. In contrast, additional cues that did not predict partner reward remained blocked from acquiring associative value. Preventing social signal exchange between the partners resulted in cues signaling partner reward remaining blocked. Taken together, these results suggest that vicarious rewards can drive reinforcement learning in rats, and that the transmission of social cues is necessary for this learning to occur.

## Introduction

Humans and other animals have developed a capacity for mutual cooperative behaviour (Nowak, 2006; Rand and Nowak, 2013; Rilling et al., 2002; Suchak et al., 2014), a preference for prosocial outcomes to familiar partners (Hernandez-Lallement et al., 2015b; Horner et al., 2011; Márquez et al., 2015) and helping behaviour towards others in need (Ben-Ami Bartal et al., 2011; Fehr and Rockenbach, 2004). These behaviours are sometimes costly, prompting questions why actor engage in them (de Waal and Suchak, 2010; Hamilton, 1963; Stevens et al., 2005; Trivers, 1971). Some researchers have focused on putative future reciprocation (Taborsky et al., 2016) as a potential driver, while others have highlighted that acting generously could generate self-reward internally (Harbaugh et al., 2007; Park et al., 2017) or through positive social reward signals such as friendly faces in humans (Spreckelmeyer et al., 2009). Indeed, the capacity to identify positive, rewarding outcomes delivered to others is a fundamental aspect of social observational learning (Zentall, 2012). Underlying some of these suggestions is the assumption that rewarding outcomes to a social partner could also represent value to oneself and thus drive a proximate reward/learning mechanism (Hernandez-Lallement et al., 2016; Ruff and Fehr, 2014). By this logic, animals, including humans, choose pro-social outcomes, cooperate or act altruistically because these actions result in vicarious reward, experienced through sensitivity to the behavioural and/or affective state of the partner (de Waal and Preston, 2017; Prochazkova and Kret, 2017), in addition to putative anticipated future reciprocal reward (Taborsky et al., 2016). One important aspect of social learning is identifying the features of the environment that predict (vicariously) rewarding outcomes, and learning the (instrumental) action sequence, appropriate to the context, for acquiring these vicariously rewarding outcomes. There is evidence that cues that predict social reward can become valuable as humans learn to respond faster to stimuli that become associated with positive social reinforcement (Jones et al., 2011) and monkeys preferred stimuli that predicted a reward delivery to a conspecific more than the stimuli that predicted no reward delivery (Chang et al., 2011). In rats, it was found that observing another rat being rewarded is (vicariously) rewarding by itself as it is accompanied by 50 kHz vocalisations, indicative of a positive appetitive state (Burgdorf et al., 2011; Panksepp, 2007), and dopamine release in the NAcc of the observer rat (Kashtelyan et al., 2014). Indeed, playback of 50 kHz leads to both an approach response (Wöhr and Schwarting, 2007) and results in dopamine release in the Nucleus Accumbens NAcc (Willuhn et al., 2014). We therefore hypothesised that vicarious reward, associated with rewards delivered to others, could also reinforce Pavlovian associative learning about novel cues, as has been found in the appetitive domain (Berridge, 2012; Schultz, 2016). To investigate our hypothesis, we use a well-established behavioural paradigm in associative learning called blocking. Kamin (1969) found, in simple stimulus-outcome association tasks, that if new stimuli are added to a stimulus that already fully predicts a reward, associative learning about those additional stimuli will be blocked. Reinforcement learning about additional stimuli can become unblocked, however, by an increase in reward value or a change in reward identity contingent on the presentation of the new stimuli. This additional or change in value is then thought to be associated to these new stimuli and thus alters their incentive value (Holland, 1984). We hypothesise that rewarding social outcomes, such as sugar pellet deliveries to a partner rat, will also be capable to unblock learning about newly added stimuli, indicative of an increased, partially vicarious value of mutual rewards relative to own-rewards. We tested this hypothesis by adopting a task from McDannald et al (2011) where unblocking is operationalised by adding additional pellet deliveries conditional on a second cue presented in compound with a learned cue that already fully predicted reward. We modified this task in such a way that the second cue is now followed by a food reward delivery to a partner rat, rather than increasing one’s own reward. In addition, a third control cue added in compound to the learned cue (on different trials) was not followed by food reward delivery to a partner rat. Concretely, we thus hypothesized that the associative learning about the second stimulus would become unblocked by mutual-reward outcomes, that is, by adding vicarious reward elicited by experiencing partner reward. In contrast, the third cue should remain blocked from acquiring associative value. We indeed found that, when tested in extinction, the unblocked second cue had acquired more associative value, as indexed by conditioned responding at the food trough, in comparison to the third blocked cue. We thus conclude that vicarious reward can support reinforcement learning processes involving novel cues. This opens up possibilities to investigate behavioural aspects of the social-value driven reinforcement learning and its associated neural basis, processes that might be disturbed in psychiatric disorders marked by impaired reinforcement learning and/or social behavior such as autism (Kohls et al., 2012) and schizophrenia (Fulford et al., 2018).Materials and

## Methods

### Subjects

56 male Long Evans rats where housed in pairs of two and kept under an inverted 12:12 h light dark cycle, in a temperature (20 ± 2 ºC) and humidity-controlled (approx. 60%) colony room. All rats had ad libitum access to food, except during the testing period. During behavioural testing, the rats where food restricted (20 grams on weekdays and 22 grams in the weekend) and maintained on a body weight of about 90% of their free-feeding weight. All testing was performed in accordance with the German Welfare Act and was approved by the local authority LANUV (Landesamt für Natur-, Umwelt und Verbraucherschutz North Rhine-Westphalia, Germany, AZ 84-02.04.2016.A522)

### Apparatus

Testing was conducted in 4 customised PhenoTyper (Noldus Information Technology) behavioural testing boxes (Fig. 1A) of 45 by 45 by 55 cm, supplemented with operant devices (Med Associates) and placed inside a custom-made sound- and lightproof ventilated box. The boxes where modified by adding a custom-made Plexiglas separation wall (Fig. 1A, left panel), which divided the box into two compartments, to allow the training of a pair of rats at the same time. The separation wall was equipped with a sliding door (dimensions: 20 by 20 cm, located at 7 cm from the left side of the Skinner box) and 4 rectangular interaction windows (Fig. 1A, left panel; size: 10 by 1.5 cm) that were positioned exactly in between the door and the wall holding the stimulation devices used for conditioning. Both compartments of the box contained a food trough **(Med Associates, ENV-254-CB)** positioned in the middle on the right side. The food troughs were adapted in such a way that the detection photobeams were positioned at the entry point of the food trough. The food trough was connected to an automated pellet dispenser (PTPD-0010, Noldus Information Technology) that delivered sucrose pellets (20 mg dustless precision pellets, Bio-Serv, Germany). Operant devices were positioned on the right side of the box at the level of the separation wall: an LED Stimulus Light (Med Associates, ENV-211m) with green cover was positioned 10 cm above the ground and a house light (Med Associates, ENV-215m) 28 cm above the ground. A speaker (Med Associates, ENV-224am) was positioned at 20 cm above the ground for the playback of auditory stimuli (Fig. 1A, right panel). Auditory stimuli were played back at a loudness of 75 dB measured with a hand-held analyser (type 2250-S from Brüel and Kjaer) right in front of the speaker. In the top cover of the Skinner boxes, a camera (Basler, acA1300-60gc, GigE) was positioned to obtain videos of the behavioural experiment at 25 fps. Analyses of the recorded videos was performed with EthoVision XT 11.5 (Noldus Information Technology). Finally, a USV-microphone was positioned next to the camera for recording ultrasonic vocalisations using Ultra Vox XT (Noldus Information Technology).

**Figure 1:**
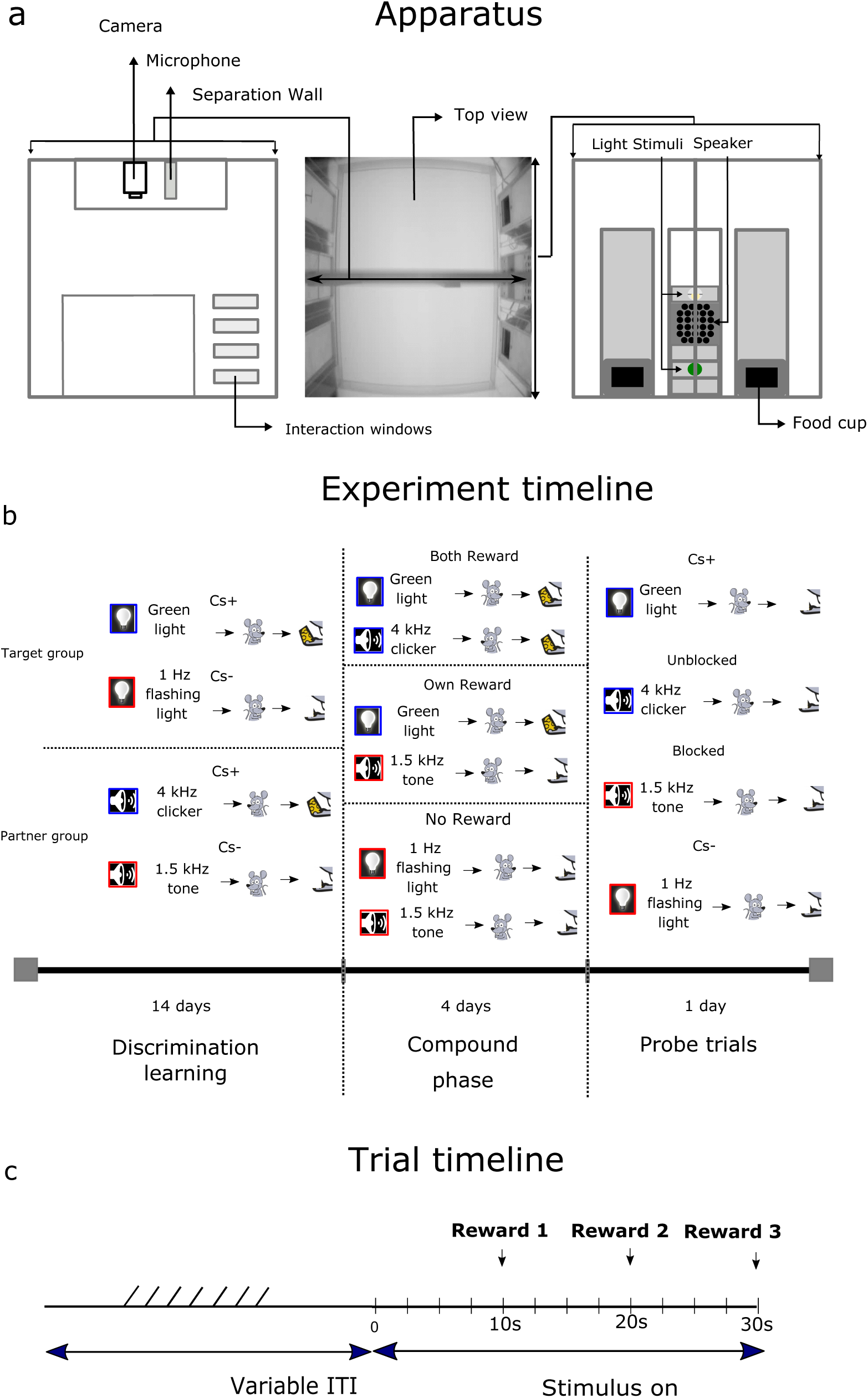
Behavioural apparatus, Experimental timeline and Trial timeline. (A) The PhenoTyper consisting of lower and upper compartment in which behavioural training took place is displayed in the middle. On the left the custom made separation wall is shown with interaction windows, camera and microphone. On the right, the right side of the PhenoTyper is displayed with the used operant devices and in both sides of the box the food cup. (B) An example experimental time line is displayed. Actor rats learn to discriminate two visual cues in upper compartment while at a different time partner rats learned to discriminate two auditory cues in the lower compartment. In the compound phase actor and partner rat are either both rewarded (Both Reward), actor rat is rewarded while the partner is not rewarded (Own Reward) or both actor and partner rat are not rewarded (No Reward). In the probe trials all learned cues are presented to the actor rat without reward. (C) Here, a timeline is shown with the different components that make up a single trial throughout the discrimination learning, compound phase and probe trials.

## Behavioral Training

### Pavlovian discrimination task

Before the start of behavioural training, rats were put on food restriction to reduce their weight to 90% of their free-feeding weight. Within a pair of cage mates, one rat was assigned at random as the actor animal, and the other as the partner animals. As a first step, they were habituated to their pre-determined training side of the customised PhenoTyper for 3 days (15 minutes per day). During this period, they could retrieve 6 pellets that were put along the edges of their respective side of the box. Subsequently, the discrimination learning phase started.

Here, the pairs of rat cage mates were divided into two groups; one group of rat pairs would learn a visual discrimination problem, and the other an auditory discrimination problem. The visual stimuli to be discriminated consisted of a houselights flashing at 1 Hz (0.1sec on, 0.9 sec off) and a steady green light; the auditory stimuli were made up by a 4.0 kHz clicker (0.1sec on, 0.9 sec off) and a 1.5 kHz (75 dB) steady tone. We counterbalanced the actor CS+ (aCS+) and actor CS- (aCS-) condition assignment to each stimulus across rat pairs. The different groups (Auditory vs Visual) were each trained alone either in the upper or lower compartment of the Skinner box, and the side assignments between actor and partner rats were counterbalanced between experiments (Fig. 1A). Each rat received 14 days of discrimination training. One daily session consisted of 40 trials, of which 20 trials were aCS+ and 20 aCS-. The order of aCS+ and aCS-trials was pseudo-randomized, with no more than 3 trials of one kind occurring in a row. Stimuli were presented for 30s and at every 10s (+ 0.1 to 0.4 sec jitter), a pellet was delivered (Fig. 1C). We trained a total of N= 20 actor rat and 20 partner rats on the discrimination problem in the experimental group and 8 actors and 8 partners in the control group. For 12 actors and 12 partners of the N=40 experimental group a second pellet dispenser was placed outside of the behavioural box, at the opposite side of where the current rat was trained, delivering pellets outside of the box. This additional dispenser placement ensured that the sound of additional pellet drops was similar to the compound conditioning phase (see below). Providing a uniform pellet delivery sound associated with self-reward pellet delivery throughout the experiment prevented any difference in pellet delivery related sounds as a source of secondary reinforcement from influencing the conditioning to added cues in the compound phase. In an additional group of 8 actor and 8 partner rats of the experimental group the second pellet dispenser was not active during the discrimination phase (Table 1). This gave us the opportunity to make direct comparison within the experimental group to investigate potential effects of secondary reinforcement (Table 1). The ITI in both experimental and control group was made up of a fixed 30s window supplemented with a randomized time window ranging from 5 to 100 seconds with steps of 5 ms, uniformly distributed. The ITIs were thus fully randomized, resulting in a total variable ITI with a mean of 80 ms. Ultrasonic vocalisations where recorded from 10s before cue onset to 20s after cue offset, for a total duration of 60s per trial. After completion of the discrimination phase, rats progressed to the compound conditioning stage.

**Table 1:** Pellet dispenser configuration and group size over different experiments.

|  | Discrimination learning | Social learning | Experimental group | Control group |
| --- | --- | --- | --- | --- |
| Experimental subgroup 1 | 1 pellet dispensers active | 2 pellet dispensers active; | 8 actors and 8 partner | 8 actors and 8 partners |
| Experimental subgroup 2 | 2 pellet dispensers active | 2 pellet dispensers active; | 12 actors and 12 partners | - |

### Compound conditioning

After discrimination training was completed, rats in the visual discrimination group received 1 day of pre-exposure to the two novel auditory stimuli while the rats in the auditory discrimination group received 1 day of pre-exposure to the two novel visual stimuli. The pre-exposure session consisted of one session with 6 trials. The stimuli were presented in a randomized order with ITIs of 15, 30 45, 60, 75 and 90s. Pre-exposure was done to reduce possible novelty-induced akinesia for each group. Cues were presented for 30s in the same manner as described for the discrimination period, only without reward. After pre-exposure, all rats received 4 days of compound training. During compound conditioning, both rats were trained together in the same Skinner box, each in their own compartment. Three different conditions were implemented; Both Reward (BR), Own Reward (OR) and No Reward (NR, Fig. 1B, middle panel). Rats from one Actor group were rewarded on BR and OR trials, but not on NR trials, while rats from the Partner group were only reward on BR trials, but not on OR and NR trials. During BR trials, both the (respective visual or auditory) CS+ of the Actor group (aCS+) and the Partner group (pCS+) were simultaneously displayed and both rats were rewarded with 3 pellets. In the OR trials, the respective aCS+ was simultaneously displayed with the aCS-of the Partner group and only the Actor group was rewarded. In the NR, both groups were presented their respective CS- (aCS-and pCS-) and none of the rats were rewarded. Assignment of visual-discrimination-trained and auditory-discrimination-trained rats to either Actor or Partner groups was counterbalanced across experiments. A compound conditioning session consisted of 20 trials per condition. The Conditions BR, OR and NR were pseudo-randomized with every condition not being repeated more than 3 times in a row. ITI randomization, stimulus presentation and reward delivery were implemented as in the discrimination phase.

### Probe trials

During probe trials, all rats were tested in isolation for one extinction session in their assigned box compartment. All stimuli were now presented in isolation, both the aCS+ and aCS-learned in the Pavlovian discrimination task as well as the two novel stimuli pCS+(Both reward CS+) and pCS- (Own Reward CS-) added in the compound phase, for which learning was hypothesised to become unblocked and blocked, respectively. Rats in both groups went through 10 trials for each of these 4 stimuli, presented in isolation and without reward delivery (Fig. 1B, right panel). The 4 stimuli were pseudo-randomized with every condition not being repeated more than 3 times in a row.

### Control experiment

In the control experiment, 8 Actor rats and 8 Partner rats went through the same three experimental conditions. The only difference here is that during the compound phase, the wall that separated the Skinner box compartments was rendered opaque by adding an additional black wall, to block contact between the Actor and Partner rats. We hypothesised that if visual, and/or auditory and/or olfactory contact between the rats facilitated the social information transmission that would unblock reinforcement learning of compound cues, then obstructing these transmission cues should impair unblocking. In the control experiment, we chose to also implement the 1-pellet dispenser condition, to bias our results to the condition where secondary reinforcement might still play a role. If differences between the control and experimental conditions would still emerge, this would strengthen the interpretation that social unblocking was driven by vicarious reward, and not secondary reinforcement learning

### Statistical data analyses

Entries into the food trough were recorded as photobeam breaks. Raw data were processed in EthoVision XT 11.5 (Noldus Information Technology) to extract our dependent variables: time spent in the food trough and number of entries in the trough (food cup rate). Food cup directed behavior in the form of time spent in the food trough and latency to entry were analysed per trial and per condition for all stages of learning; further analysis and graph preparation was performed using custom-made scripts in MATLAB (version 2014b, MathWorks). All statistics was performed using SPSS (IBM Corp. Released 2017. IBM SPSS Statistics for Windows, Version 25.0. Armonk, NY: IBM Corp). To assess the strength of learning during discrimination and compound conditioning, only the first 10s of the cue period was analysed to avoid the influence of reward delivery/omission feedback (McDannald et al., 2011) and the time spent in the food cup was used as measure for conditioned responding. In the probe trials however reward was absent, therefore here we analysed both 10s and 30s period. Previously (Burke et al., 2008; McDannald et al., 2011) used the percentage of time spent at the food trough and the food cup rate to assess value unblocking and identity unblocking, respectively as measures for conditioned responding. As social unblocking is thought to mainly reflect value unblocking, we report the percentage of time spent as our main outcome parameter. For completion, we also report food cup rate to assess identity unblocking. Discrimination learning performance was quantified by averaging responding to the cues over the last 4 days of training and comparing the mean between aCS+ and aCS- and difference scores of CS+ - CS-for contrasting cue modalities using paired sample t-tests. Performance in the compound phase was quantified using a 2 factor repeated measures ANOVA on the mean response rate per day across conditions (BR, OR and NR) and post hoc tests were performed to assess the significance of any differences between conditions, corrected for multiple comparisons. Simple contrasts were calculated to further compare the difference between conditions on the different days against the last day of compound conditioning. Performance in the probe trials was assessed by averaging responding of the actor rats time spent in the food cup over 5 bins (2 trials per bin) and running a two factor repeated measures ANOVA over these bins and the 4 stimuli types (aCS+, pCS+ (unblocked), pCS- (blocked) and aCS-) separately for experimental and control experiments. Differences between conditions and bins were assessed with post-hoc tests, again corrected for multiple comparisons. Here, simple contrasts were calculated, comparing the difference between conditions on the different bins against the last bin of extinction. A non-parametric Friedman two-way ANOVA test on the differences among the latency to entries was furthermore performed including all trials (excluding all non-entered trials) between the conditions in the probe trials. Differences between conditions were assessed with post-hoc tests, again corrected for multiple comparisons. For the direct comparison between Experiment and control group in the probe trials we performed a mixed repeated measures ANOVA with factors trials (trial 1 to 6) and stimuli (aCS+, pCS+ (unblocked), pCS- (blocked) and aCS-) and experiment group as a between subjects factor (Experiment, Control). Differences between conditions between experiments were assessed with post-hoc tests, corrected for multiple comparisons. Finally, to further in depth look at the difference between experiment and control group difference scores were calculated for every available contrast (aCS+/aCS-, pCS+/pCS-, pCS+/aCS- and pCS-/aCS) and for these contrast a two factor repeated measures anova was calculated. For all RM anovas Mauchly’s test of sphericity was performed and only when significant the Greenhouse-Geisser correction was applied.

## Results

All groups of actor and partner rats were initially trained separately on a Pavlovian discrimination problem. Subsequently, the rats went through a social learning phase were actor rats could learn to associate additional compounded cues with different reward outcomes delivered to the partner rat (social unblocking). Finally, we tested the associative strength of all cues, each presented in isolation, in a probe phase without a reward. In a control experiment, we impeded the exchange of visual information by implementing an opaque wall. In the following sections, we show the actor rats conditioned responses with the time spent in the food cup and latency to entry as dependent variables for the three stages of learning per group (experimental, control) and show a comparative analyses of experimental versus control group. Subsequently, we look at food cup rate in the last stage of learning with similar comparative analyses of experimental versus control. Finally, we show the moderating effect of potential secondary reinforcement due to additional pellet deliveries on the strength of social unblocking.

### Experimental group results

#### Discrimination learning

Actor rats (N=20) were trained on a discrimination task with counterbalanced visual or auditory exemplars. All actor rats developed a conditioned response to their own aCS+, resulting in an increase in time spent in the food trough on aCS+ trials in anticipation of reward, independent of cue modality or identity. Concurrently, they learned to expect no reward during aCS-presentations, as witnessed by a steady decrease in time spent in the food trough on aCS-trials (Fig. 2A). A paired samples *t*-test examining the mean responding over the last 4 days of conditioning was performed. We found a significant difference in time spent in the food trough between the aCS+ (M = 58.76, SD=12,86) and aCS- (M=21.19, SD= 13,21; *t*(19) = 12.109, p < 0.001 (Fig. 2A). We furthermore found that when looking at the difference scores of CS+/CS-that the discrimination for visual cues did not differ in comparison to auditory cues (M = -3.55, SD=21.42; *t*(8) = 0.497, p < 0.632). These results indicate that our rats discriminated between auditory cues of 1.5 kHz vs. 4 kHz clicker (0.1 Ms per 1s on) and between visual green vs. white flicker (0.1 Ms per 1s on) and that that was no difference between modalities.

**Figure 2.**
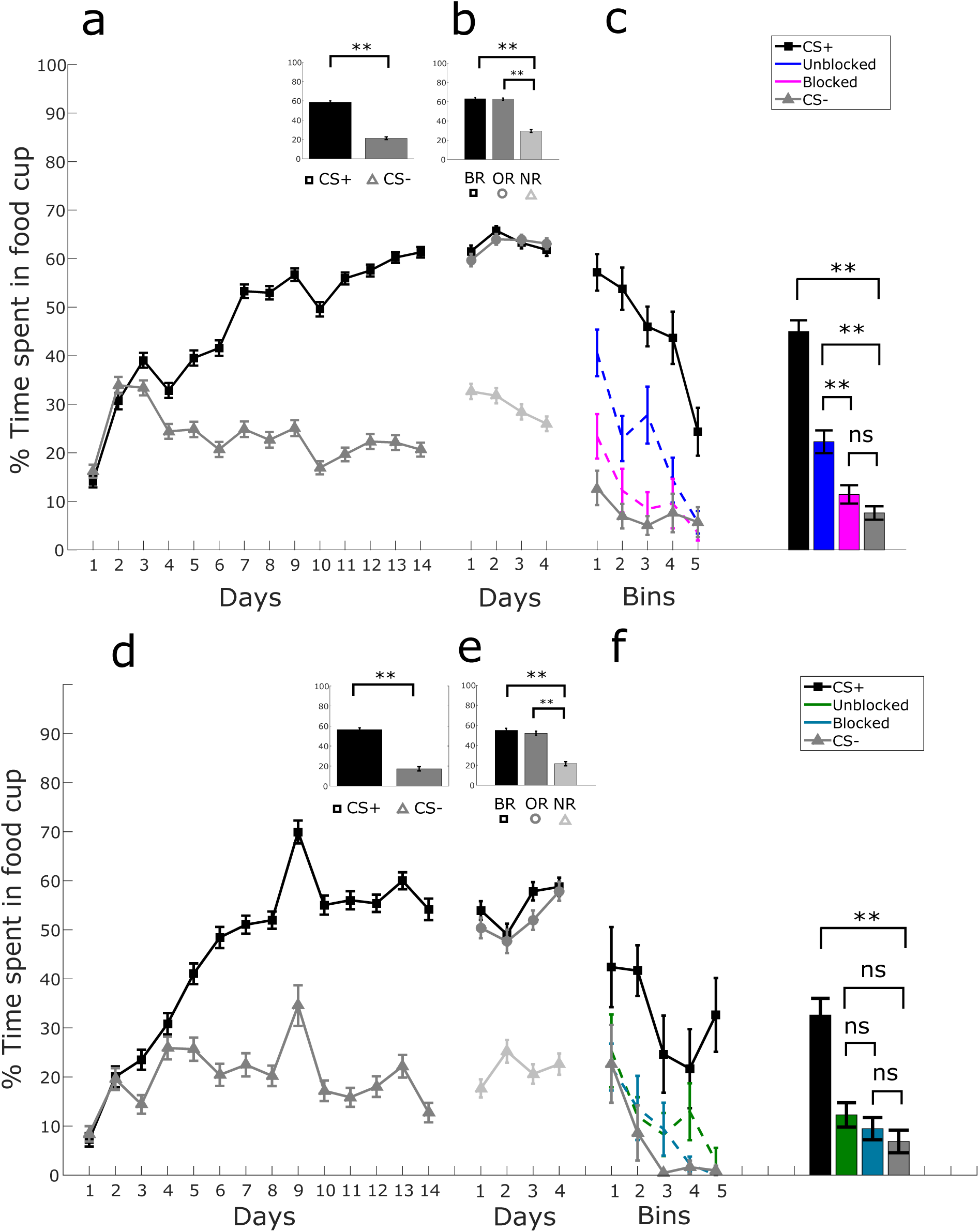
Conditioning per experimental phase. Experimental group. (A) Percentage of time spent in food cup for discrimination learning between aCS+ and aCS- over days. (B) Percentage of time spent in food cup for the compounds BR (aCS+, pCS+), OR (aCS+, pCS-) and NR (aCS-,pCS-) over days. (C) Percentage of time spent in the food cup during the probe trials over 10 trials. Averaged time spent in the food cup over 10 probe trials between aCS+, pCS+ (unblocked), pCS- (blocked) and aCS-. Control group. (D) Percentage of time spent in food cup for discrimination learning between aCS+ and aCS- over days. (E) Percentage of time spent in food cup for the compounds BR (aCS+,pCS+), OR (aCS+, pCS-) and NR (aCS-,pCS-) over days. (F) Percentage of time spent in the food cup during the probe trials over 10 trials. Averaged time spent in the food cup over 10 probe trials between aCS+, pCS+(unblocked), pCS- (blocked) and aCS-. Error bars indicate SEM. (*p < 0.05; **p < 0.01,***p < 0.001)

#### Social learning

In this phase, rats were trained together. The aCS+/aCS-of the actor and pCS+/pCS-of the partner were combined in three compound combinations with the following reward outcomes: Both Reward (BR), Own Reward (OR) and No Reward (NR; Fig. 1B). We chose to omit the condition where the target rats would *not* receive reward, while the partner rats would, to avoid a potential reward/value conflict due to disadvantageous inequity aversion (Fehr and Schmidt, 1999), which has been reported in rats as well (Oberliessen et al., 2016). Rats’ conditioned responses to these compound cues are shown and a direct comparison of these responses to the original aCS+ and aCS-cues was made (Fig. 1B), both indexed by time spent in the food cup and the food cup rate. In the subsequent analysis, only the behavior of the Actor rats is reported. A 2-factor repeated measures ANOVA with condition and day as factors and the time spent in food trough as dependent variable was performed. We found a significant main effect of condition type (F _(2, 38)_ = 417,04, p < 0.01, η_p_^2^ = 0.956). As expected, this effect was explained by a greater responding on BR and OR trials (including the aCS+) compared to NR trials (including the aCS-; both p < 0.01, Bonferroni-corrected; Fig. 2B) and no difference between responding on the BR compared to the OR condition (p = 1.00). Furthermore, a paired samples *t*-test revealed that responding on BR (M = 61.8, SD = 7.08) or OR trials (M = 63.1, SD = 6.93) did not significantly differ from the average response rate over the last 4 days of discrimination learning for the aCS+ (M = 58.76, SD = 12.85,21; *t*(19) = 0.907, p = 0.376 and *t*(19)=1.334, p = 0.198). We found no main effect of day (F_(3, 57)_ = 2.248, p = 0.092, η_p_^2^ = 0.106) but did find a significant interaction between condition and day (F_(6,114)_ = 2.795, p=0.014, η_p_^2^ = 0.128). Simple contrasts revealed that the difference between condition BR and NR (F_(1,19)_ = 7,238, p = 0.014) and the difference between conditions OR and NR (F_(1,19)_ = 6.898, p = 0.017) was significantly higher for day 4 compared to day 1 (Fig. 2B). The increase in differences between BR and NR trials and between OR and NR trials, when contrasting day 1 and day 4, is driven by a decrease of responding on the NR trials over days, as witnessed by a significant decrease when directly comparing day 1 (M = 32.63, SD= 8.47) to day 4 (M = 25.97, SD= 7.68; *t*(19) = 2.898, p = 0.009; Fig. 2B, light grey line). A paired samples *t*-test revealed that responding on NR trials was significantly higher on day 1 (M = 32.64, SD = 8.47) than responding, averaged over the last 4 days of discrimination learning, for aCS-trials (M = 21.18, SD = 13.21); *t*(19)= 3.218, p = 0.005, suggesting perhaps a temporary enhanced motivational effect of the partner presence in the NR compound impairing previously learned inhibitory motor responses related to the aCS-. Responding on NR day 4 however (M = 25.97, SD = 7,68;) did not differ anymore from responding to aCS-averaged over the last 4 days of discrimination learning; *t*(19)= 1.523, p = 0.114. We conclude from these results that adding a second cue, which predicts either a reward (BR) or no reward (OR) to the partner, to the actor rats aCS+ did not change the actor rat responding on BR and OR in comparison to its originally learned aCS+ response at the end of discrimination. However, adding a second cue, which predicts no reward (NR) to the partner, on a learned aCS-response did change the actor rats’ responding on the NR condition. Here, a transient increase in responding was found that subsided over 4 days of compound training. Most importantly, both BR and OR trials, including the aCS+ as part of the compound, still elicited more responses than NR (aCS-compound) trials.

### Probing vicarious reward value

In the probe trials, we aimed to show the effect of associative learning driven by self and vicarious reward. In an extinction setting, the cues were presented in isolation, omitting reward. Associative value of each cue was indexed by the time spent in the food cup, the food cup rate and the latency to entry. We show the percentage conditioned responding of the actor rats to the first 10 seconds of 10 presentations each of the aCS+ and aCS-, the pCS+ (unblocked) cue associated with an added reward to the partner (BR) and the pCS- (blocked) cue associated with no reward to the partner (OR) and no reward to self (NR). A two-factor repeated measures ANOVA with stimulus type and bin as factors and the time spent in the food trough as the dependent variable was performed. Summary statistics (F-stats, p-values, effect sizes) can be found in the supplemental materials, Table 1. We found a significant main effect of probe trial condition on time spent in food trough (F _(3, 57)_ = 83.287, p < 0.01, η_p_^2^ = 0.814). As expected, time spent in food trough was higher for aCS+ than aCS-trials (Mean Difference = 37.406, Std Error = 2.964, p < 0.001; Cohen’s d = 1.92, Fig. 2C). Critically, pairwise comparisons revealed that the actors time spent in food trough was also higher for pCS+ cues than for pCS-cues across the 5 bins of extinction (Mean Difference = 10.822, Std Error = 2.598, p = 0.003, Cohen’s d = 0.53; Fig. 2C). Furthermore, we found a significantly higher responding to the pCS+ cue compared to the aCS-cue (Mean Difference = 14.646, Std Error = 2.245, p < 0.001, Cohen’s d = 0.78) while responding to the pCS-cue was not significantly different from the aCS-cue (Mean Difference = 3.824, Std Error = 1.915, p = 0.362, Cohen’s d = 0.24; Fig. 2C). Additionally, we found a main effect of bin number on time spent in food trough (F _(4, 76)_ = 18.809, p < 0.01, η_p_^2^ = 0.497), reflecting the extinction process, and finally, we found an interaction between condition type and bin number on the time spent in food trough (F _(12,228)_ = 2.930, p = 0.001, η_p_^2^ = 0.134). Simple contrasts revealed that the difference between pCS+ and pCS-was significantly higher in bin 1 (F_(1,19)_ = 4.94, p = 0.039) and bin 3 (F(_1,19)_ = 7.127, p = 0.015), and trending for bin 2 (F_(1,19)_ = 3.365, p = 0.082) in comparison to that difference observed in bin 5. Simple contrasts furthermore revealed that the difference in responding for the pCS+ and aCS-cues was significantly higher in bin 1 (F_(1,19)_ = 42.316, p = 0.000), bin 2 (F_(1,19)_ = 10.857, p = 0.004) and bin 3 (F_(1,19)_ = 15.530, p = 0.001) than the difference observed in bin 5). A non-parametric Friedman two-way ANOVA test on the differences in the latency to entry for all trials (excluding all non-entered trials) between the conditions aCS+ (median = 3.32), pCS+ (median = 3.56), pCS- (median = 6.8) and aCS- (median = 9.58) during extinction was performed. We found a significant effect of probe trial condition on the latency to entry (X^2^_(3, N = 62)_ = 23.733, p < 0.001). Post hoc comparisons revealed that latencies to entry were significantly shorter for aCS+ than both pCS- (p < 0.001) and aCS- (p < 0.001) but not the pCS+ cue (p = 0.203). However, pCS+ entry latencies did not differ significantly from pCS- (p = 0.309) and aCS- (p = 0.203); pCS- and aCS-entries also did not differ significantly (p = 1.000). These results suggest that the latencies to entry for the pCS+ cue fall in between the aCS+ and the pCS-/aCS-cues (Figure S2C).

Taken together, these results show that the actor rats exhibited more food cup directed behavior for the pCS+ cue than both the aCS- and pCS-cue over 10 trials of extinction. This means that when actor rats have fully learned a stimulus-reward association producing reward for themselves, adding a cue that predicted an additional reward delivery to a partner rat unblocked associative learning about this cue, possibly due to vicarious reward experience. We conclude from the contrast analyses explaining the interaction effect that the unblocking of the novel cue lasts for approximately 6 trials and will use this analysis window going forward. In contrast, rats did not spent more time in the food cup for the pCS-cue than the aCS-, suggesting that additional cues that did not predict vicarious reward remained blocked from acquiring associative value.

### Control group

In this control experiment, we impeded the exchange of direct social contact by introducing an opaque wall that blocked visual contact, covered the interaction windows (Fig. 1) and dampened auditory cues from the partner compartment. All other task parameters were exactly the same (see Table 1).

#### Discrimination learning

We replicated the discrimination learning results from the experimental group showing that rats again developed a conditioned response to the aCS+, but not to the aCS- (Fig. 2D). A paired sampled t-test revealed a significant difference in time spent in food trough between the aCS+ (M = 56.40, SD =12.89) and aCS-(M =17.20, SD = 10,18); *t*(7) =11.54, p < 0.001 (Fig. 2D). We furthermore also found that when looking at the difference scores of CS+/CS-that the discrimination for visual cues did not differ in comparison to auditory cues (M = 12.53, SD= 8.63; *t*(8) = 2.904, p = 0.062). These results indicate that our rats here again discriminated between auditory cues of 1.5 kHz vs. 4 kHz clicker (0.1 Ms per 1s on) and between visual green vs. white flicker (0.1 Ms per 1s on) and that there was no difference between modalities. Finally, control rats also developed a shorter latency to entry for the aCS+ and a larger latency to entry for the aCS-over training (Figure S2D).

#### Social learning

In the compound phase, rats were again trained side by side on the compound cues associated with the social outcomes Both Reward (BR), Own Reward (OR) and No Reward (NR; Fig. 1B). This time, however, we aimed to block visual and auditory information exchange between the partners. We hypothesized that blocking social exchange would prevent the experience of vicarious reward, and thus would prevent social unblocking. A 2-factor repeated measures ANOVA with condition and day as factors and the time spent in food trough as dependent variable was performed. Mauchly’s test of sphericity was significant for both the main effect of the factor condition and days. Therefore, for these effects, the Greenhouse-Geisser corrected values are reported. As under the experimental conditions, we found a significant effect of condition type on time spent in food trough (F_(1.175, 8.228)_ = 46.407, p < 0.01, η_p_^2^ = 0.869). This effect was again explained by a greater responding on BR and OR trials compared to NR trials (p = 0.01, Fig. 2E). We found no effect of days on time spent in the food trough (F_(1.294, 9.059)_ = 1.501, p = 0,262, η_p_^2^ = 0.177) but again did find an interaction between day and condition (F_(6, 42)_ = 2.873, p= 0.019, η_p_^2^ = 0.291)). We first compared conditioned responding on the last 4 days of conditioning with the first day of BR and OR. A paired samples *t*-test revealed that responding for BR trials did not differ on day 1 (M = 58.45, SD = 9.40) in comparison to the average of the last 4 days of responding for the CS+ (M=56.40, SD= 12.89); *t*(7)=0.398, p = 0.702) and when comparing OR (M = 50.38, SD= 15.84) with the CS+ (M=56.40, SD= 12.89); *t*(7)=-1.779, p = 0.118). To explore further the meaning of the interaction between day and condition we looked at contrast scores. Contrasts revealed that the difference between condition BR and NR (F(1,7) = 13.043, p = 0.009) and the difference between conditions OR and NR (F(1,7) = 7.00, p = 0.033) was significantly higher for day 4 compared to day 2 but found no difference between day 1 and 4. The increase in difference between the BR and NR and difference between OR and NR when contrasting day 2 and day 4 was not driven by a decrease in responding to the NR from day 1 (M = 17.68, SD = 7.56) to day 4 (M = 22.63, SD = 10.59; *t*(7)= -1.041, p = 0.333) (Fig 2E) as was the case in the experimental group results (Fig 2B). Paired samples *t*-test revealed furthermore that the time spent in the food cup for NR on day 1 (M = 17.68, SD = 7.56) was not significantly higher than responding on the last 4 days of CS- (M = 17.20, SD = 10.17); *t*(7) = 0.137, p = 0.895). These results indicate that adding a second cue that predicts either reward or no reward to the partner when social information exchange is impeded does not influence time spent in the nose poke for a self-reward. Both BR and OR trials, including the CS+ as part of the compound, still elicited a higher time spent in the food cup than NR (CS-compound) trials. However when social information exchange was blocked, we found that observing a partner not being rewarded did not result in an increase in the time spent in the food cup for the NR on day 1. NR value is comparable to CS-value throughout compound conditioning as indexed by the time spent in the food cup and thus no further modulation through learning processes became apparent.

#### Probing vicarious reward value

In the control experiment probe trials, we aimed to show the effect of impeding social information exchange on vicarious reward experience through unblocking. We again show the conditioned responding of the actor rats to 10 presentations each of all cues. A two-factor repeated measures with condition and bin as factors and the time spent in the food cup as dependent variable was performed. Summary statistics (F-stats, p-values, effect sizes) can be found in the supplemental materials, Table 1. As in the experimental condition, we found a significant main effect of probe trial condition type on time spent in food trough (F_(3, 21)_ = 15.327, p < 0.01, η_p_^2^ = 0.686). We furthermore found a main effect of bin number on time spent in food trough (F_(1.540, 10.781)_ = 18.809, p < 0.01, η_p_^2^ = 0.509). We did not find an interaction between condition type and bin number on the time spent in food trough (F_(12,84)_ = 1.225, p = 0.280, η_p_^2^ = 0.149). Performing post hoc comparisons, we again found that responding to the aCS+ was higher than the aCS- (Mean Difference = 32.615, Std Error = 6.019, p = 0.021, Cohen’s d = 1.45). Crucially however, in contrast to the experimental condition we found that the actors time spent in the food trough was not significantly different between pCS+ (unblocked) and pCS- (blocked) cues across the 5 bins of extinction (Mean Difference = 2.80, Std Error = 2.662, p = 1.000, Cohen’s d = 0.19; Fig. 2F). Moreover, in this control experiment, time spent in the food trough for both the pCS+ cue and the pCS-cue was not different from the aCS- (Mean Difference = 5.415; 2.615, Std Error = 2.662; 2.247; p= 0.622; p=1.00, Cohen’s d = 0.37, 0.19 respectively).

These results show that, contrary to the findings for the experimental group, the rats in the control experiment did not show more food cup directed behavior for the pCS+ cue than both the aCS- and blocked cue over 10 trials of extinction. This suggests that when actor rats have fully learned a stimulus-reward association producing reward for themselves, adding a cue that predicted an additional reward delivery to a partner rat does not unblock associative learning when social information exchange and putative vicarious reward experience is blocked.

#### Direct comparison between Experimental and Control conditions

To examine the strength of the unblocking effect with respect to the control condition, we first applied a mixed repeated measures ANOVA design with trial type (aCS+, pCS+ (unblocked), pCS- (blocked), aCS-) and trial 1-6 (bin 1-3) as within subject factors and group (experimental (N=20) vs control N= 8) as between subject factor with time spent in the food cup the first 10 second after the cue onset as dependent variable. Summary statistics (F-stats, p-values, effect sizes) can be found in the supplemental materials, Table 1. We found a significant main effect of trial type (F_(3, 78)_ = 52.790, p < 0.001, η_p_^2^ = 0.670), an interaction effect of Experiment * trial type (F_(3,78)_ = 5.464, p= 0.002, η_p_^2^ = 0.174) and an effect of trial (F_(5,130)_ = 8.446, p< 0.001, η_p_^2^ = 0.245). Post-hoc comparison reveals that the actors’ responding to the pCS+ cue differs significantly from the pCS-cue (Mean Difference = 15.753, Std Error = 3.188, p < 0.001, Cohen’s d = 0.57) in the experimental group but not the control group (mean difference = 0.175, std error = 5.073, p = 1.00, Cohen’s d = 0.01; Fig 3A). The aCS+, however, differs significantly from the aCS- (mean difference = 44.10, std error = 3.514, p < 0.001, Cohen’s d = 2.86) in the experimental group as well as in the control group (mean difference = 25.667, std error = 5.556, p = 0.001, Cohen’s d = 1.53). To examine this contrast in findings between the experimental and control conditions more in depth we calculated difference scores for the direct comparison of the aCS+/aCS-, pCS+/pCS-, pCS+/aCS- and pCS-/aCS-contrasts between groups. We examined the difference scores of the pCS+/pCS-contrast with a two way repeated measures ANOVA with group (Experimental, control) and trial (1-6) as factors. We found no significant within-subject main effect of trial number F_(5, 130)_ = 0.715, p = 0.613 η_p_^2^ = 0.027) and no interaction effect (F_(5,130)_ = 0.554, p= 0.735, η_p_^2^ = 0.021). However, a significant main (between subject) effect of group (F_(1, 26)_ = 6.823, p = 0.013, η_p_^2^ = 0.208; Fig. 3B) revealed that the percent difference in responding between pCS+ and pCS-cues was higher for the experimental group than the control group (mean difference = 15.578, std error = 5.964). We similarly examined the difference scores for the pCS+/aCS-contrast with a two way repeated measures ANOVA. We found no significant (within subject) main effect of trial on group (F_(5,130)_ = 0.589, p = 0.708) and no interaction effect (F_(5,130)_ = 1.750, p= 0.128). But here as well, we found a significant main between subject effect of group (F_(1, 26)_ = 8.614, p =0.007, η_p_^2^ = 0.249 Fig. 3C) when comparing the pCS+ / aCS-contrast score (mean difference = 17.567, std error = 5.985). As expected, when comparing responding for the pCS- / aCS-contrast (mean difference = 1.988, std error = 4.234), we found no significant (within subject) main effect of trial (F_(5,130)_ = 1.425, p = 0.219) and no interaction effect (F_(5,130)_ = 0.597, p= 0.702). Crucially we also found no significant main between subject effect of group (F_(1, 26)_ = 0.221, p =0.643, η_p_^2^ = 0.008; Fig. 3E), suggesting that the ratio of responding to the blocked and learned CS-is not influenced by the addition of the blackout wall. We then examined the difference scores for the aCS+/aCS-contrast with a two way repeated measures ANOVA. We found no significant main (within subject) effect of trial on group (F_(5,130)_ = 1.133, p = 0.346) and no interaction effect (F_(5,130)_ = 0.604, p= 0.697). We did however find an unexpected significant main (between subject) effect of group (F_(1, 26)_ = 7.862, p =0.009, η_p_^2^ = 0.232; Fig. 3D) when comparing the aCS+/aCS-difference score (mean difference = 18.433, std error = 6.574) suggesting some between-group differences in the efficacy of aCS+ conditioning. Finally, we also investigated the unblocking effect on food cup occupancy by looking at the time spent during the whole 30s cue-on period. As the animals would expect the first reward delivery (on some trials) after 10 seconds, this analysis necessarily is influenced by the animals’ response to these omissions during the probe phase. For this analysis, we again used a mixed repeated measures ANOVA design with trial type (aCS+, pCS+ (unblocked), pCS- (blocked), aCS-) and trial 1-6 (bin 1-3) as within subject factors and group (experimental (N=20) vs control N= 8). A similar pattern as for the 10s data held for examining the full 30 seconds after the cue onset as dependent variable. We again found a significant main effect of trial type (F_(3, 78)_ = 186.988, p < 0.001, η_p_^2^ = 0.878); applying the greenhouse-geisser correction we found an interaction effect of Experiment * trial type (F_(2.280,59.284)_ = 6.251, p= 0.002, η_p_^2^ = 0.194) and an effect of bin (F_(2,52)_ = 28.267, p< 0.001, η_p_^2^ = 0.521). Post-hoc comparison revealed here as well that the pCS+ cue differed significantly from the pCS-cue (Mean Difference = 8.950, std error = 2.606, p =0.012, Cohen’s d = 0.57) in the experimental group but not the control group (Mean Difference = -1.237, std error = 4.120, p = 1.00, Cohen’s d = -0.10; Fig 3a - s1). The aCS+ differed significantly from the aCS- (mean difference = 56.44, std error = 2.254, p < 0.001, Cohen’s d = 4.02) in the experimental group and in the control group (mean difference = 39.831, std error = 3.434, p < 0.001, Cohen’s d = 2.31). To examine this contrast in findings between the experimental and control conditions more in depth we calculated difference scores for the direct comparison of the aCS+/aCS-, pCS+/pCS-, pCS+/aCS- and pCS- /aCS-contrasts between groups. We examined the difference scores of the contrast with a two way repeated measures ANOVA with group (Experimental, control) and trial (1-6) as factors. For the pCS+/pCS-contrast we found no significant within-subject main effect of trial number (F_(5, 130)_ = 1.061, p = 0.385, η_p_^2^ = 0.039) and no interaction effect (F_(5,130)_ = 1.097, p= 0.365, η_p_^2^ = 0.040). However, a significant main (between subject) effect of group (F_(1, 26)_ = 4.631, p = 0.041, η_p_^2^ = 0.151; Fig. 3B - s1) revealed that the percent difference in responding between pCS+ and pCS-cues was higher for the experimental group than the control group (mean difference = 9.554, std error = 4.435). For the aCS+/aCS-contrast we found a significant within-subject main effect of trial number (F_(5,130)_ = 2.633, p = 0.0.092, η_p_^2^ = 0.092) and no interaction effect (F_(5,130)_ = 0.944, p= 0.455, η_p_^2^ = 0.035). However, a significant main (between subject) effect of group (F_(1, 26)_ = 15.518, p = 0.001, η_p_^2^ = 0.374; Fig. 3D - s1) revealed that the percent difference in responding between aCS+ and aCS-cues was higher for the experimental group than the control group (mean difference = 16.609, std error = 4.216). For the pCS+/aCS-contrast we found a significant within-subject main effect of trial number (F_(5, 130)_ = 2.735, p = 0.022, η_p_^2^ = 0.095) and an interaction effect (F_(5,130)_ = 4.927, p < 0.001, η_p_^2^ = 0.159). A significant main (between subject) effect of group (F_(1, 26)_ = 4.631, p = 0.041, η_p_^2^ = 0.151; Fig. 3C-s1) revealed that the percent difference in responding between pCS+ and aCS-cues was higher for the experimental group than the control group (mean difference = 12.412, std error = 4.422). For the pCS-/aCS-contrast we found no significant within-subject main effect of trial number (F_(3.662, 95.203)_ = 1.299, p = 0.277, η_p_^2^ = 0.048) and no interaction effect (F_p (3.662,95.203) p_ = 2.290, p = 0.071, η_p_^2^ = 0.081). Crucially, we did not find a significant main (between subject) effect of group (F_(1, 26)_ = 0.786, p = 0.384, η_p_^2^ = 0.029; Fig. 3E-s1) revealed that the percent difference in responding between pCS- and aCS-cues was not higher for the experimental group than the control group (mean difference = 3.083, std error = 3.479).

**Figure 3.**
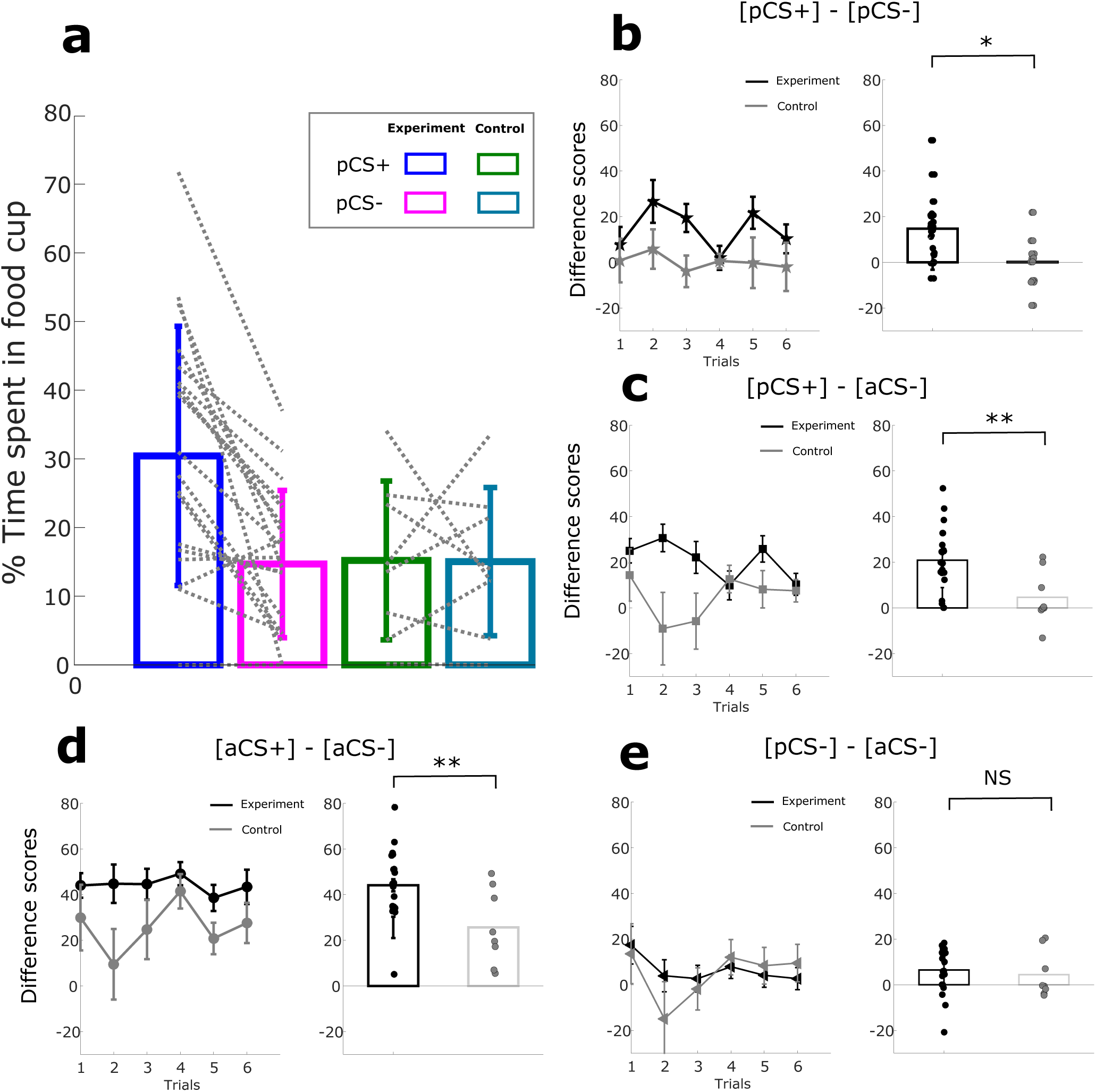
Food cup occupancy in Experimental and control group. (A) Mean percentage of Time spent in the food cup during the first 10 seconds after cue onset for experiment vs control for trial 1 to 6. (B) Difference scores of the percentage of time spent in food cup over 6 trials for the [pCS+] - [pCS-] difference scores. Bar plots show average over 6 trials of the [pCS+] - [pCS-] difference scores with dots showing the mean per rat (C) Difference of the percentage of time spent in food cup over 6 trials for the [pCS+] - [Cs-] difference scores. Bar plots show average over 6 trials of the [pCS+] - [Cs-] difference scores with dots showing the mean per rat (D) Difference of the percentage of time spent in the food cup over 6 trials for the [Cs+] - [CS-] difference scores. Bar plots show average over 6 trials of the [Cs+] - [CS-] difference scores with dots showing the mean per rat. (E) Difference of percentage of the time spent in food cup responding over 6 trials for the [pCS-] - [Cs-] difference scores. Bar plots show average over 6 trials of the [pCS-] - [Cs-] difference scores with dots showing the mean per rat. Line graph error bars indicate the SEM. (*p < 0.05; **p < 0.01; ***p < 0.001)

We conclude from these results that the compounded cue pCS+ has become unblocked for the actor rat in the experimental group, but not the control group, as witnessed by a significantly larger unblocked vs. blocked contrast in the experimental vs. control group in both the first 10 seconds after cue onset and the whole 30 seconds period. We attribute this differential social unblocking effect to the difference in experienced vicarious reward, which was presumably attenuated by impeding social information exchange in the control group. In both groups, the compounded cue pCS-remained blocked from acquiring associative value (in comparison to the aCS-), as would be expected from learning theory. Though we found a magnitude difference in the aCS+/aCS-contrast between experimental and control groups, we note that the aCS+/aCS-contrast was significant by itself in both groups as well, while this was not the case for the pCS+/pCS-contrast.

### Food cup rate

Burke et al., 2008 found that changing the sensory identity (flavour) of an outcome associated with an added cue in a compound also unblocked this cue and that this identity unblocking was captured by scoring the food cup rate, e.g. the frequency or number of entries into the food cup irrespective of the total duration of visits. In our paradigm, social unblocking could also be interpreted as a reward identity switch in that the additional partner outcome changes the sensory aspects reward by virtue of the partner receiving and eating the rewards. Food cup rate in the probe trials, next to food cup occupancy, could therefore potentially reflect model-based reward identity unblocking and therefore could provide insight in the influence of sensory features of social unblocking. Alternatively, if the additional partner reward is interpreted solely as a change in value, but not identity, we would hypothesize that food cup rate would not be affected.

#### Experimental group

We performed a two factor repeated measures with condition and bin as factors and the food cup rate as dependent variable. We found a significant main effect of probe trial condition type on food cup rate (F_(1.676, 31.853)_ = 24.940, η_p_^2^ = 0.568, p < 0.01). Pairwise comparisons revealed that the food cup rate was descriptively higher across the 5 bins of extinction in response to pCS+ cues than to pCS-cues (p = 0.084) and significantly higher than to CS-cues (p< 0.01, Fig. S1C). In contrast, responding to the pCS-cue did not differ from responding to the CS- (p = 0.190). We furthermore found a main effect of bin number on food cup rate (F_(4, 76)_ = 18.629, η_p_^2^ = 0.528, p < 0.01) as well, in line with extinction processes and did not find a interaction between condition type and bin number on the food cup rate (F_(3.953,75.116)_ = 1.630, η_p_^2^ = 0.079,p = 0.176).

#### Control group

Here, we also performed a two factor repeated measures with condition and bin as factors and the food cup rate as dependent variable. We found a significant main effect of probe trial condition type on food cup rate (F_(1.292, 9.042)_ = 10.979, p = 0.007). We furthermore find a main effect of bin number on time spent in food trough (F_(4, 28)_ = 10.335, p < 0.001). Finally, we found no interaction between condition type and bin number on the time spent in food trough (F_(12, 84)_ = 1.229, p = 0.235). Pairwise comparisons revealed that the food cup rate was marginally higher in CS+ cues compared to CS-cues across the 5 bins of extinction (p = 0.057, Fig. S1F). We furthermore found that the food cup rate was not different in pCS+ cues in comparison to pCS-cues across the 5 bins of extinction (p = 1.000, Fig. 3C). Finally, here both the pCS+ cue and pCS-cue were not different then the CSmin (p = 0.330, p= 0.190)

#### Direct comparison between Experimental and Control conditions

To further explore the food cup rate as a measure of the potential identity unblocking effect in the experimental vs. control condition, we applied a mixed repeated measures ANOVA design with trial type (aCS+, pCS+ (unblocked), pCS- (blocked), aCS-) and bin 1-3 as within subject factors and group (experimental (N=20) vs control N= 8) as between subject factor with food cup rate in the first 10 second after the cue onset as dependent variable. We found a significant main effect of trial type (F_(1.808, 47.017)_ = 16,696, p < 0.001, η_p_^2^ = 0.395) and an effect of trial (F_(2,52)_ = 19.671, p< 0.001, η_p_^2^ = 0.431). We did not find an interaction effect of Experiment * trial type (F_(3,78)_ = 0.637, p= 0.593, η_p_^2^ = 0.024).

It is clear from these results that the food cup rate is not significantly higher for pCS+ cue in comparison to the pCS-cue in both the experimental group and control group. There appears to be some trend for an increased responding for the pCS+ compared to the pCS- and CSmin in the experimental group. We can conclude however, that identity unblocking in the form of an increased food cup rate, does not seem to play a major role in social unblocking.

#### Control for secondary reinforcement

Besides the vicarious experience of reward, other confounding factors could have contributed to learning/unblocking in our paradigm. Most notably, sources of secondary reinforcement should be excluded as potential drivers of learning. During discrimination learning, the actor rat is conditioned to receive pellets contingent on its aCS+. Afterwards, in the compound phase, the rat is presented with an auditory– visual compound. Instead of one pellet drop (self-reward), now, on some trials, two pellets drop simultaneously (Both Reward trials). It is possible that the additional pellet delivery sound acted as a third CS+ in the compound, besides the aCS+ and pCS+. Because the sound of the pellet dispenser is already associated with the aCS+ of the actor rat, it is possible that the appetitive value increased with the intensity of this cue (two pellets dropping instead of one), thus enhancing the total value of cue configuration, leading to unblocking of the pCS+. To control for this possible source of secondary reinforcement, in a subgroup of rats, we added a pellet dispenser aimed outside the box already during the discrimination phase, providing the same acoustic features of pellet delivery to the target rat, without presenting additional reward. Here, we compare results between these conditions by again looking at responding during the first 10 seconds of the probe phase, which would equate to time period in which one extra pellet would be delivered. We performed a mixed repeated measures ANOVA design with trial type (pCS+, pCS-) and trial 1 to 6 as within subject factors and group (experimental_1: N=8; one pellet added vs. experimental_2: N=12; no new pellets added vs. Control: N= 8; one pellet added) as a between-subjects factor (see also Table 1). We found a significant main effect (F_(1, 25)_ = 21.546, p < 0.001, η_p_^2^ = 0.463) of trial type, an interaction of Group * trial type (F_(2,25)_ = 7.396, = 0.003, η_p_^2^ = 0.372) and an effect of trial number (F_(2,50)_ = 5.456, p < 0.001, η_p_ ^2^ = 0.179). Post hoc comparison reveals that the pCS+ cue differs significantly from the pCS-cue group 1 (one pellet added; mean difference = 24.808, std error = 4.578, p < 0.001, Cohen’s d = 1.13), group 2 (no new pellets added; mean difference = 9.717, std error = 3.738, p = 0.015, Cohen’s d = 0.50) but not the control group (one pellet added; mean difference = 0.175, std error = 4.578, p = 0.970, Cohen’s d = 0.01; Fig 4).

**Figure 4.**
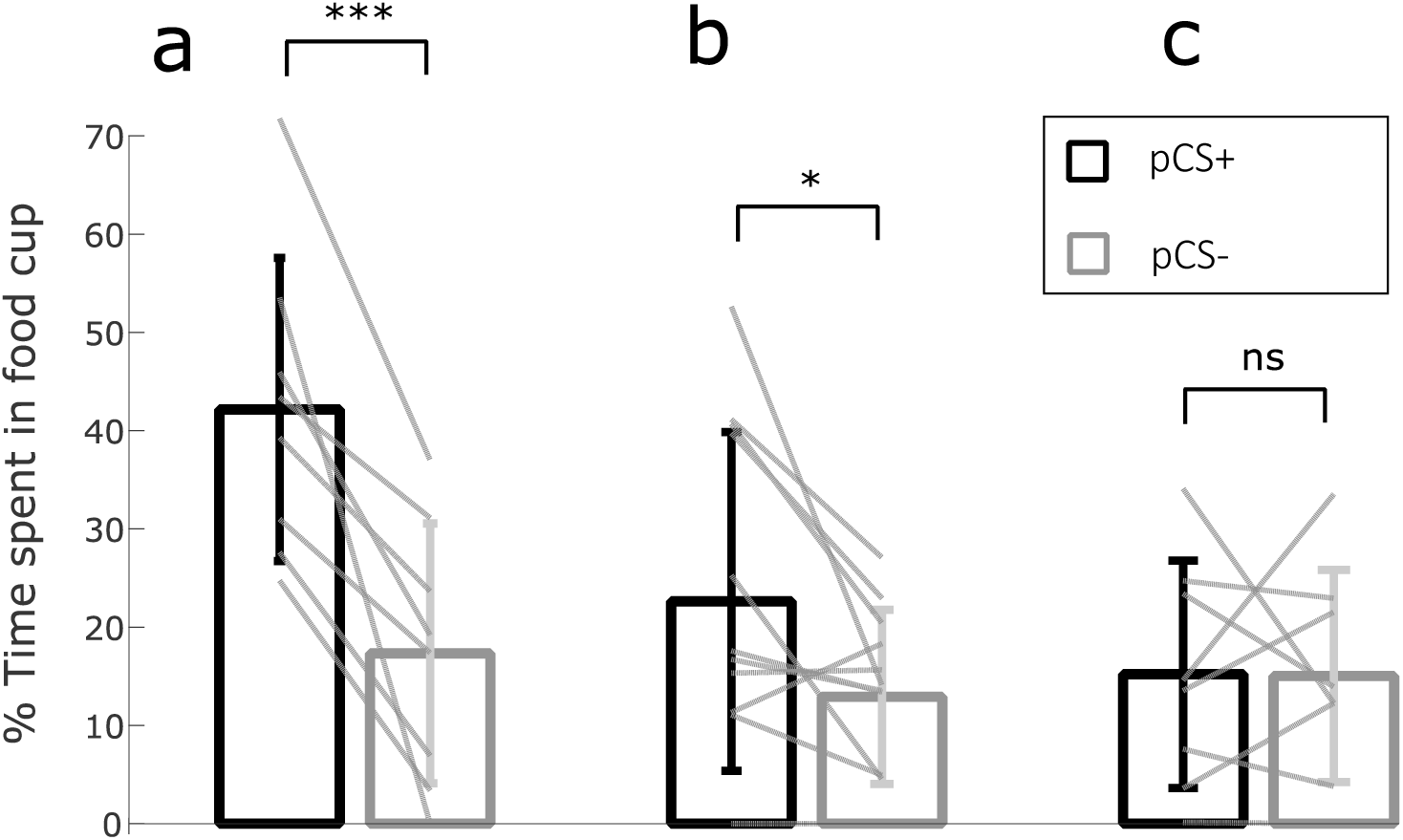
Effect of secondary reinforcement on food cup occupancy in the first 10 seconds after cue onset. (A) Mean percentage of spent in the food cup for trial 1 to 6 for the experimental subgroup 1 (N =8) without additional pellet dispensers present during discrimination learning. (B) Mean percentage of time spent in the food cup for trial 1 to 6 for the experimental subgroup 2 (N=12) with 1 additional pellet dispensers present during discrimination learning. (C) Mean percentage of time spent in the food cup for the control group without additional pellet dispensers during discrimination

Finally, we compared results between these conditions by looking at the probe phase’s first full cue on period which would equate to the addition of 3 extra pellets. We performed a mixed repeated measures ANOVA design with trial type (pCS+, pCS-) and bin (1 - 3) as within subject factors and 3 group (experimental_1 (n=8); 3 pellet added) vs experimental_2 (n=12; no new pellets added), Control N= 8; 3 pellet added) factors as between subject factor. We find a significant main effect (F_(1, 25)_ = 9.246, p = 0.005, η_p_^2^ = 0.270), an interaction effect of Group * trial type (F_(2,25)_ = 5.218, p = 0.013, η_p_^2^ = 0.294) and an effect of bin (F_p (2,50) p_ = 6.975, p = 0.002, η_p_^2^ = 0.002). Post hoc comparison reveals that the pCS+ cue differs significantly from the pCS-cue in group 1 (one pellet added; mean difference = 15.775, std error = 3.814, p < 0.001, Cohen’s d = 1.04), but not in group 2 (no new pellets added; mean difference = 4.399, std error = 3.114, p = 0.170, Cohen’s d = 0.33) but not the control group (mean difference = -1.273, std error = 3.814, p = 0.971, Cohen’s d = -0.10).

While descriptively, the magnitude of the unblocking effect is larger when not controlling for additional pellet drops (Experimental_1) than when such a control is implemented (Experimental_2), we conclude that the unblocking effect still exist when explicitly controlling for additional pellets falling in the compound phase.

## Discussion

### Summary

Social valuation is crucial in forming and maintaining social relationships and, presumably, in experiencing the pleasurable and reinforcing aspects of social interaction. However, it remained unclear whether vicarious reward value, which we define here as value derived from social signals associated with reward to another (Ruff and Fehr, 2014), could drive learning just as self-experienced value. If this would be the case, then vicariously experienced reward should be able to reinforce behaviour in a formal Pavlovian learning paradigm. Here we addressed this question by introducing a novel social unblocking task. We find that vicarious reward experience, operationalized in this task as rewards delivered to social partners, can indeed drive learning about novel stimuli. After having fully learned that a specific cue predicted a self-reward, learning about a second cue delivered in compound was blocked, as predicted by learning theory (Fanselow and Wassum, 2016; Rescorla and Wagner, 1972), when no additional self or other reward was contingent on this cue. Learning was unblocked, however, by providing an additional reward delivered to the partner simultaneously with the fully predicted self-reward. Finally, preventing the exchange of social information in the compound learning phase impeded unblocking of the novel stimuli. These results suggest that vicarious reward experience can indeed drive learning processes, in line with formal behavioural learning theory.

### Learning Theory

Our results extend previous work by Peter Holland (Holland, 1984) and Geoffrey Schoenbaum (Lopatina et al., 2015; McDannald et al., 2011) on unblocking in appetitive Pavlovian conditioning. These authors found that rats, after learning that distinct cues have specific food outcomes, can show unblocking of learning for cues added in compound, when self-rewards were altered by increasing reward value (e.g. an upward shift from 1 to 3 pellets) or a change in reward identity (same reward type but a shift in reward features such as flavour). In contrast, learning was blocked when no such reward change occurred (e.g. same reward amount or same identity). According to reinforcement learning theory, the upshift or change in identity led to a discrepancy between the expected reward (1 pellet) and the received reward (3 pellets), thus producing a reward prediction error. The theory states that if the added cue reliably predicts the increase in reward/ identity outcome, it will acquire the value inherent in the reward itself (Sutton and Barto, 1981). The main indicator of learning about the value of a (novel) cue in our social unblocking paradigm is an increase in the time spent in the food cup in the probe (extinction) phase (Lopatina et al., 2015; McDannald et al., 2011). Indeed, we observed an increase in the time spent in the food cup for the unblocked cue, predicting mutual rewards to the actor rat and its conspecific, relative to the blocked cue, predicting own-rewards only, when presenting the cue in isolation without reward delivery.

From this result, we conclude that the added novel cue has acquired value through vicarious reward inherent in the “Both Reward” delivery by way of reinforcement learning mechanisms. Analogous to the description above, our result mimics food cup responding that represent the appetitive US-directed ‘wanting’ response (Berridge, 2012). This indicates that the incentive salience of the unblocked cue can, in addition to being changed by appetitive reward, also be changed by vicarious reward experience. Further research is necessary to see whether cues associated with vicarious reward or social reinforcement can also act as a conditioned reinforcer for instrumental responses of rats, as has been found humans (Lehner et al., 2017), in a similar way as has been found for appetitive cues (Burke et al., 2008; Kruse et al., 1983; Rescorla, 1994). Finally, it is important to investigate if cues predicting vicarious rewards can guide rats’ choices in a social setting. It has been found that rats choose a reward arm in a T-maze that leads to play behavior more than an arm leading to a social encounter where play was absent (Humphreys and Einon, 1981). Furthermore, social play can induce a social place preference (Calcagnetti and Schechter, 1992) and rats are willing to lever press for social play reinforcement (Achterberg et al., 2016). Our task indicates that the unblocked cue has attained the rewarding properties of social reward and it is therefore likely that a presented unblocked cue would be preferred over a blocked cue when tested in a choice task. Our findings that unblocking can be driven by vicarious reward experience as well as Pavlovian appetitive value adds to the emerging literature showing that animals attach value to rewards delivered to conspecifics (Hernandez-Lallement et al., 2016; Kashtelyan et al., 2014) and learn about cues that predict rewards delivered to others. Monkeys, for example, prefer cues that are associated with reward delivery to another monkey over cues that were associated with juice delivery to a chair with no monkey in it and this preference was absent, in the non-social condition, when there was only a juice bottle present (Chang et al., 2011). By contrast, not all unequal reward structures lead to positive social value presumably motivating pro-social behavior. A recent study shows that macaques increase licking frequency in line with a higher probability of self-reward but decrease their anticipatory licking with increased probability of reward delivery to another monkey (Noritake et al., 2018). The authors interpret this decrease of anticipatory licking as an indicator of the negative affect associated with unequal disadvantageous reward pay-outs. Both monkeys and rats have been found to have a distaste for these unequal pay-outs (Brosnan and De Waal, 2003; Oberliessen et al., 2016). It is thus clear that monkeys can learn to associate cues with both negative and positive outcomes delivered to others. Our results provide evidence that, next to monkeys, rats as well can learn about cues that are associated with reward delivered to others. The social reinforcement learning hypothesis (Hernandez-Lallement et al., 2016) proposes that integration of social signals expressed by partners can aid in making appropriate decisions in a social context. Evidence for this hypothesis comes from the prosocial choice task (PCT) in which it was found that, rewards delivered to oneself and to a partner are preferred over a reward delivery only to the actor himself in both monkeys (Horner et al., 2011) and rats (Hernandez-Lallement et al., 2015a; Márquez et al., 2015). In rats, it was found that this effect was modulated by the behavior displayed by the other rat (Márquez et al., 2015) and that this effect was impaired when the partner was replaced by a toy (Hernandez-Lallement et al., 2015b) or when the display of the partner’s preference was impeded. Adding to these results, we found that learning occurs when additional reward was delivered to the partner and that preventing the exchange of social information by an opaque wall impeded this effect. These results also fit the cascade model of representation of cognitive (social) structures that predicts partial contributions that sum up over multiple modalities of social signals (Kalenscher et al, in press). We thus expect that associative learning of the actor rats in our task is driven by a similar social reinforcement learning process, dependent on the transmission of social signals between the actor and partner rat (Nicol, 1995). These signals could manifest in the following aspects; confounding factors, auditory transfer, visual observation and olfactory/tactile social interaction. In the following paragraph, these potential modes of social information transfer will be discussed.

### Social Information Transfer

#### 1. Auditory transfer of affective state/value

Our data show that the possibility for social information exchange was necessary to produce the social unblocking effect. One possibility is that social information was transferred through auditory transfer of affective state/value between rats. It has been found that rats observing conspecifics being rewarded produce 50 kHz vocalisations and show an increase in dopamine release (Kashtelyan et al., 2014) and Playback of 50 kHz vocalisation furthermore motivates rats to approach the location of playback and increase their USV production (Wöhr and Schwarting, 2007). Importantly, during playback of 50 kHz calls, dopamine release in the Nucleus Accumbens is transiently increased (Willuhn et al., 2014). Prosocial choices in rats, as found by Hernandez-Lallement in the PCT (Hernandez-Lallement et al., 2015a), are therefore likely to be driven by communicative social reinforcement signals (Hernandez-Lallement et al., 2016), such as 50 kHz vocalisations expressed by the partner rat. Recently, it has been found that supressing VTA dopamine transients by means of optogenetics during the compound phase of a non-social blocking task impairs unblocking of novel cue value and features (Chang et al., 2017). Furthermore, increasing VTA dopamine activity can drive learning about novel cue-reward associations in an unblocking paradigm (Keiflin et al., 2019). Taken together, these findings suggest that unblocking in our task could occur through social reinforcement by way of 50 kHz calls-associated dopamine release. Some evidence that 50 kHz can enhance the value of novel cues comes from a study by Saito et al (2016) that found that the playback of 50 kHz ultrasonic vocalisations, 20 min. prior to hearing a neutral tone, increased lever presses of rats during subsequent presentation of the neutral tone more so than in a control condition where there was no playback. The amount of lever presses became more similar to the number of lever presses observed in response to a previously conditioned rewarding stimulus (positive bias), indicating that the playback of 50 kHz USVs induced a positive affective state that transferred onto the neutral stimuli and thereby enhancing its value. In our task, positive reinforcement of the partner can be perceived as both surprising and rewarding to the actor rat and this could therefore increase the partner’s and/or actor’s 50 kHz vocalisations. Future work will indicate whether the production of 50 kHz USVs (unpublished observations) during the BR trials in the compound phase can indeed contribute to dopamine release in the actor rat and thus drive learning i.e. unblocking of the additional compound cue that is associated with partner reward.

#### 2. Visual observational learning

Another possibility is that social information was transferred through visual observational learning. Previous experiments in rats have shown that, during learning, animals can develop both goal tracking towards the location of the US and sign tracking behavior showing orienting and enhanced responding to the location of the CS (Cleland and Davey, 1983). This enhanced responding is indicative of an enhanced incentive salience of the cue. We observed that some animals in our task indeed showed coordinated cue-directed behavior during the compound phase. This conforms to the definition of stimulus enhancement in social learning: when observation of an action leads the observer to increase the proportion of its behavior directed towards the location or object of the demonstrator’s (Spence, 1937)). Such coordinated putative sign-tracking could therefore facilitate learning in the actor rats about the cues inducing cue-directed behavior in the partner rat and presumably increase sign tracking behavior for the actor rat towards the unblocked cue. As an example of such stimulus enhancement, Heyes et al. (Heyes et al., 2000) showed that rats observing other rats performing a lever pressing discrimination task have a higher lever-pressing rate for a previously observed ACS+ lever. In our task, responding at the food cup by actor rats might be enhanced through observational conditioning, i.e. by observing the conditioned approach of the partner on BR trials in contrast to OR/NR trials. Our experimental setup was not designed to gauge the influence of observational learning on unblocking because both animals approach the reward site simultaneously during BR conditions. Furthermore, the observed latency to entry was not different between conditions in the compound phase and probe phase. This implies though that a pure matching response is unlikely, but leaves room for a role for coordinated sign tracking during the both reward cue and subsequent observation of the partner’s approach (on BR trials) or lack thereof (on OR/NR trials). Trial by trial computational modelling of the influence of the partner rat’s sign tracking behavior and approach behavior on the actor rat’s actions could lead to further insights regarding the putative role of stimulus enhancement and observational learning in this task.

#### 3. Olfactory and tactile information exchange

Finally, social information exchange could have occurred through olfactory and tactile information exchange. It has been found that in rats, food preferences can be transferred from demonstrators to observers and that for this effect to occur, the scent of the food together with the presence of carbon disulphide (CS2) on the breath of the demonstrator are necessary (Galef, Jr., 2001). Preliminary observations of the interaction between our rats in this task showed that facial touch occurs throughout all conditions in the interaction window (Fig. 1B). The presence of pellet consumption related olfactory cues and associated CS2 on the partner’s breath could thus have aided the actor rats in identifying BR trials, while an absence or decrease in these indicator smells might indicate OR/NR trials. In addition, it has been found that facial touch is associated with an increase in the production of 50 kHz calls (Rao et al., 2014).

Within a BR condition, where sniffing of dyadic olfactory cues might be enhanced, social face touch could potentially further enhance the production of 50 kHz calls and thus contribute to unblocking.

## Conclusion

Overall, these data provide evidence that vicarious reward experience can drive reinforcement learning in rats, and that the transmission of social cues is necessary for this learning. Further experiments should be conducted to reveal which mode(s) of social information processing are necessary and sufficient to drive unblocking through social value. Overall, our novel behavioural paradigm could be used to further explore how rats learn about value in social contexts and is well suited to probe the neural circuits involved in social reinforcement learning.

## Author Contributions

### Conceptualization

MvW and SvG; Methodology: MvW, SvG and JH; Formal Analysis: SvG; Investigation: SvG and JH; Resources: MvW and TK; Writing – Original Draft: SvG; Writing – Review & Editing: MvW, TK and SvG; Visualization; SvG; Supervision, MvW; Funding Acquisition: MvW; Project Administration: SvG, MvW.

## References

Achterberg EM, Wm Van Kerkhof L, Servadio M, Mh Van Swieten M, Houwing DJ, Aalderink M, Driel N V, Trezza V, Vanderschuren LJ. 2016. Contrasting Roles of Dopamine and Noradrenaline in the Motivational Properties of Social Play Behavior in Rats. Neuropsychopharmacology 41212:858–868. doi:10.1038/npp.2015.212

Ben-Ami Bartal I, Decety J, Mason P. 2011. Empathy and pro-social behavior in rats. Science 334:1427–30. doi:10.1126/science.1210789

Berridge KC. 2012. From prediction error to incentive salience: mesolimbic computation of reward motivation. Eur J Neurosci 35:1124–43. doi:10.1111/j.1460-9568.2012.07990.x

Brosnan SF, De Waal FB. 2003. Monkeys reject unequal pay. Nature 425:297–299.

Burgdorf J, Panksepp J, Moskal JR. 2011. Neuroscience and Biobehavioral Reviews Frequency-modulated 50 kHz ultrasonic vocalizations : a tool for uncovering the molecular substrates of positive affect. Neurosci Biobehav Rev 35:1831–1836. doi:10.1016/j.neubiorev.2010.11.011

Burke KA, Franz TM, Miller DN, Schoenbaum G. 2008. The role of the orbitofrontal cortex in the pursuit of happiness and more specific rewards. Nature 454:340–4. doi:10.1038/nature06993

Calcagnetti DJ, Schechter MD. 1992. Place conditioning reveals the rewarding aspect of social interaction in juvenile rats. Physiol Behav 51:667–672. doi:10.1016/0031-9384(92)90101-7

Chang CY, Gardner M, Di Tillio MG, Schoenbaum G. 2017. Optogenetic Blockade of Dopamine Transients Prevents Learning Induced by Changes in Reward Features. Curr Biol 27:3480-3486.e3. doi:10.1016/j.cub.2017.09.049

Chang SWC, Winecoff AA, Platt ML. 2011. Vicarious reinforcement in rhesus macaques (macaca mulatta). Front Neurosci 5:27. doi:10.3389/fnins.2011.00027

Cleland GG, Davey GCL. 1983. Autoshaping in the rat: The effects of localizable visual and auditory signals for food. J Exp Anal Behav 40:47–56. doi:10.1901/jeab.1983.40-47

de Waal FB, Suchak M. 2010. Prosocial primates: selfish and unselfish motivations. Philos Trans R Soc L B Biol Sci 365:2711–2722. doi:365/1553/2711 [pii]10.1098/rstb.2010.0119

de Waal FBM, Preston SD. 2017. Mammalian empathy: behavioural manifestations and neural basis. Nat Rev Neurosci 18:498–509. doi:10.1038/nrn.2017.72

Fanselow MS, Wassum KM. 2016. The origins and organization of vertebrate pavlovian conditioning. Cold Spring Harb Perspect Biol 8:1–28. doi:10.1101/cshperspect.a021717

Fehr E, Rockenbach B. 2004. Human altruism: economic, neural, and evolutionary perspectives. Curr Opin Neurobiol 14:784–90. doi:10.1016/j.conb.2004.10.007

Fehr E, Schmidt KM. 1999. A theory of fairness, competition and cooperation. Q J Econ 114:817–868.

Fulford D, Campellone T, Gard DE. 2018. Social motivation in schizophrenia: How research on basic reward processes informs and limits our understanding. Clin Psychol Rev 63:12–24. doi:10.1016/j.cpr.2018.05.007

Galef, Jr. B. 2001. Social Influences on Food Choices of Norway Rats and Mate Choices of Japanese Quail. Int J Comp Psychol 14. doi:https://escholarship.org/uc/item/4fw7m0dk

Hamilton WD. 1963. The Evolution of Altruistic Behavior. Am Nat 97:354.

Harbaugh WT, Mayr U, Burghart DR. 2007. Neural Responses to Taxation and Voluntary Giving Reveal Motives for Charitable Donations. Science (80-) 316:1622–1625. doi:10.1126/science.1140738

Hernandez-Lallement J, van Wingerden M, Marx C, Srejic M, Kalenscher T. 2015a. Rats prefer mutual rewards in a prosocial choice task. Front Neurosci 8. doi:10.3389/fnins.2014.00443

Hernandez-Lallement J, Van Wingerden M, Marx C, Srejic M, Kalenscher T. 2015b. Rats prefer mutual rewards in a prosocial choice task. Front Neurosci 9:1–9. doi:10.3389/fnins.2014.00443

Hernandez-Lallement J, van Wingerden M, Schäble S, Kalenscher T. 2016. A Social Reinforcement Learning Hypothesis of Mutual Reward Preferences in RatsCurrent Topics in Behavioral Neurosciences. pp. 159–176. doi:10.1007/7854_2016_436

Heyes CM, Ray ED, Mitchell CJ, Nokes T. 2000. Stimulus Enhancement: Controls for Social Facilitation and Local Enhancement. Learn Motiv 31:83–98. doi:10.1006/lmot.1999.1041

Holland PC. 1984. Unblocking in Pavlovian appetitive conditioning. J Exp Psychol Anim Behav Process 10:476.

Horner V, Carter JD, Suchak M, de Waal FBM, Waal FBM De. 2011. Spontaneous prosocial choice by chimpanzees. Proc Natl Acad Sci U S A 108:13847–51. doi:10.1073/pnas.1111088108

Humphreys AP, Einon DF. 1981. Play as a reinforcer for maze-learning in juvenile rats. Anim Behav 29:259–270. doi:10.1016/S0003-3472(81)80173-X

Jones RM, Somerville LH, Li J, Ruberry EJ, Libby V, Glover G, Voss HU, Ballon DJ, Casey BJ. 2011. Behavioral and neural properties of social reinforcement learning. J Neurosci 31:13039–45. doi:10.1523/JNEUROSCI.2972-11.2011

Kamin LJ. 1969. Predictability, surprise, attention and conditioning. B A Campbell, R M Church Punishm Aversive Behav 279–296.

Kashtelyan V, Lichtenberg NT, Chen ML, Cheer JF, Roesch MR. 2014. Observation of reward delivery to a conspecific modulates dopamine release in ventral striatum. Curr Biol 24:2564–2568. doi:10.1016/j.cub.2014.09.016

Keiflin R, Pribut HJ, Shah NB, Janak PH, Keiflin R, Pribut HJ, Shah NB, Janak PH. 2019. Ventral Tegmental Dopamine Neurons Participate in Article Ventral Tegmental Dopamine Neurons Participate in Reward Identity Predictions 93–103. doi:10.1016/j.cub.2018.11.050

Kohls G, Chevallier C, Troiani V, Schultz RT. 2012. Social “wanting” dysfunction in autism: neurobiological underpinnings and treatment implications. J Neurodev Disord 4:10. doi:10.1186/1866-1955-4-10

Kruse JM, Overmier JB, Konz WA, Rokke E. 1983. Pavlovian conditioned stimulus effects upon instrumental choice behavior are reinforcer specific. Learn Motiv 14:165–181. doi:10.1016/0023-9690(83)90004-8

Lehner R, Balsters JH, Herger A, Hare TA, Wenderoth N. 2017. Monetary, Food, and Social Rewards Induce Similar Pavlovian-to-Instrumental Transfer Effects. Front Behav Neurosci 10:1–12. doi:10.3389/fnbeh.2016.00247

Lopatina N, McDannald MA, Steyer C V, Sadacca BF, Cheer JF, Schoenbaum G. 2015. Lateral orbitofrontal neurons acquire responses to upshifted, downshifted, or blocked cues during unblocking. Elife 4:e11299. doi:10.7554/eLife.11299

Márquez C, Rennie SM, Costa DF, Moita MA. 2015. Prosocial Choice in Rats Depends on Food-Seeking Behavior Displayed by Recipients. Curr Biol 25:1736–1745. doi:10.1016/j.cub.2015.05.018

McDannald MA, Lucantonio F, Burke KA, Niv Y, Schoenbaum G. 2011. Ventral striatum and orbitofrontal cortex are both required for model-based, but not model-free, reinforcement learning. J Neurosci 31:2700–5. doi:10.1523/JNEUROSCI.5499-10.2011

Nicol CJ. 1995. The social transmission of information and behaviour. Appl Anim Behav Sci 44:79–98. doi:10.1016/0168-1591(95)00607-T

Noritake A, Ninomiya T, Isoda M. 2018. Social reward monitoring and valuation in the macaque brain. Nat Neurosci 21:1452–1462. doi:10.1038/s41593-018-0229-7

Nowak MA. 2006. Five rules for the evolution of cooperation. Science (80-) 314:1560–1563. doi:10.1126/science.1133755

Oberliessen L, Hernandez-Lallement J, Schäble S, van Wingerden M, Seinstra M, Kalenscher T. 2016. Inequity aversion in rats, Rattus norvegicus. Anim Behav 115:157–166. doi:10.1016/j.anbehav.2016.03.007

Panksepp J. 2007. Neuroevolutionary sources of laughter and social joy: modeling primal human laughter in laboratory rats. Behav Brain Res 182:231–44. doi:10.1016/j.bbr.2007.02.015

Park SQ, Kahnt T, Dogan A, Strang S, Fehr E, Tobler PN. 2017. A neural link between generosity and happiness. Nat Commun 8:15964. doi:10.1038/ncomms15964

Prochazkova E, Kret ME. 2017. Connecting minds and sharing emotions through mimicry: A neurocognitive model of emotional contagion. Neurosci Biobehav Rev 80:99–114. doi:10.1016/j.neubiorev.2017.05.013

Rand DG, Nowak MA. 2013. Human cooperation. Trends Cogn Sci 17:413–425. doi:10.1016/j.tics.2013.06.003

Rao RP, Mielke F, Bobrov E, Brecht M. 2014. Vocalization–whisking coordination and multisensory integration of social signals in rat auditory cortex. Elife 3. doi:10.7554/eLife.03185

Rescorla RA. 1994. Control of Instrumental Performance by Pavlovian and Instrumental Stimuli. J Exp Psychol Anim Behav Process 20:44–50. doi:10.1037/0097-7403.20.1.44

Rescorla, Wagner. 1972. Theory of Classical Conditioning.

Rilling J, Gutman D, Zeh T, Pagnoni G, Berns G, Kilts C. 2002. A neural basis for social cooperation. Neuron 35:395–405. doi:S0896627302007559 [pii]

Ruff CC, Fehr E. 2014. The neurobiology of rewards and values in social decision making. Nat Rev Neurosci 15:549–562. doi:10.1038/nrn3776

Saito Y, Yuki S, Seki Y, Kagawa H, Okanoya K. 2016. Cognitive bias in rats evoked by ultrasonic vocalizations suggests emotional contagion. Behav Processes 132:5–11. doi:10.1016/j.beproc.2016.08.005

Schultz W. 2016. Dopamine reward prediction-error signalling: a two-component response. Nat Rev Neurosci 17:183–95. doi:10.1038/nrn.2015.26

Spence KW. 1937. Experimental studies of learning and the higher mental processes in infra-human primates. Psychol Bull 34:806–850. doi:10.1037/h0061498

Spreckelmeyer KN, Krach S, Kohls G, Rademacher L, Irmak A, Konrad K, Kircher T, Gründer G. 2009. Anticipation of monetary and social reward differently activates mesolimbic brain structures in men and women. Soc Cogn Affect Neurosci 4:158–65. doi:10.1093/scan/nsn051

Stevens JR, Cushman F a., Hauser MD. 2005. Evolving the Psychological Mechanisms for Cooperation. Annu Rev Ecol Evol Syst 36:499–518. doi:10.1146/annurev.ecolsys.36.113004.083814

Suchak M, Eppley TM, Campbell MW, de Waal FBM. 2014. Ape duos and trios: Spontaneous cooperation with free partner choice in chimpanzees. PeerJ 2014:1–19. doi:10.7717/peerj.417

Sutton RS, Barto AG. 1981. Toward a modern theory of adaptive networks: expectation and prediction. Psychol Rev 88:135–70.

Taborsky M, Frommen JG, Riehl C. 2016. Correlated pay-offs are key to cooperation. Philos Trans R Soc B Biol Sci 371:20150084. doi:10.1098/rstb.2015.0084

Trivers RL. 1971. The evolution of reciprocal altruism. Q Rev Biol 46:35–57. doi:doi:10.1086/406755

Willuhn I, Tose A, Wanat MJ, Hart AS, Hollon NG, Phillips PEM, Schwarting RKW, Wöhr M. 2014. Phasic dopamine release in the nucleus accumbens in response to pro-social 50 kHz ultrasonic vocalizations in rats. J Neurosci 34:10616–23. doi:10.1523/JNEUROSCI.1060-14.2014

Wöhr M, Schwarting RKW. 2007. Ultrasonic communication in rats: can playback of 50-kHz calls induce approach behavior? PLoS One 2:e1365. doi:10.1371/journal.pone.0001365

Zentall TR. 2012. Perspectives on observational learning in animals. J Comp Psychol 126:114–128. doi:10.1037/a0025381

