## Supplemental Materials for "Vicarious reward unblocks associative learning about novel cues in male rats"

| *Table S1. Probing the associative value of vicarious reward outcomes* | | | | |
|---|---|---|---|---|
| ***Experimental group*** | F value **(main effect)** | P value | $\eta_p^2$ | |
| RM anova (Within Subject; aCS+,aCS-, pCS+, pCS-; bin 1- 5) | $_{(3, 57)}$ = 83.287 | **p < 0.01** | $\eta_p^2$ = 0.814 | |
| Post hoc comparisons for main effect | **Mean difference** | **Std error** | **P value** | **Cohen's d** |
| aCS+ \| aCS- | 37.406 | 2.964 | **< 0.001** | 1.92 |
| pCS+ \| aCS- | 14.646 | 2.245 | **< 0.001** | 0.78 |
| pCS+ \| pCS- | 10.822 | 2.598 | **0.003** | 0.53 |
| pCS- \| aCS- | 3.824 | 1.915 | 0.362 | 0.24 |
| ***Control group*** | **F value** (main effect) | **P value** | $\eta_p^2$ | |
| RM anova (Within Subject; aCS+,aCS-, pCS+, pCS-; bin 1- 5) | $_{(3, 21)}$ = 15.327 | < 0.01 | | |
| Post hoc comparisons for main effect | **Mean difference** | **Std error** | **P value** | **Cohen's d** |
| aCS+ \| aCS- | 32.615 | 6.019 | **0.021** | 1.45 |
| pCS+ \| aCS- | 5.415 | 2.662 | 0.622 | 0.37 |
| pCS+ \| pCS- | 2.80 | 2.662 | 1.000 | 0.19 |
| pCS- \| aCS- | 2.615 | 2.247 | 1.000 | 0.19 |
| ***Experimental - Control Group comparison*** | **F value** (interaction effect) | **P value** | $\eta_p^2$ | |
| Mixed RM anova (Within Subject; aCS+,aCS-, pCS+, pCS-; trial 1 - 6 & Between Subjects; Experimental vs control) | $_{(3,78)}$ = 5.464 | 0.002 | 0.174 | |
| Post hoc comparisons for interaction effect | **Mean difference** | **Std error** | **P value** | **Cohen's d** |
| **Experimental group** | | | | |
| aCS+ \| aCS- | 44.10 | 3.514 | **< 0.001** | 2.86 |
| pCS+ \| pCS- | 15.753 | 3.188 | **< 0.001** | 0.57 |
| **Control group** | | | | |
| aCS+ \| aCS- | 25.667 | 5.556 | **0.001** | 1.53 |
| pCS+ \| pCS- | 0.175 | 5.073 | 1.00 | 0.01 |
| ***Experimental - Control Group; Difference scores*** | | | | |
| RM anova (Within Subject; aCS+/aCS-, pCS+/pCS-, pCS+/aCS- and pCS-/aCS- contrasts, trial 1 – 6; Between Subjects; Experimental vs control) | **F value** (Between subjects effect) | **P value** | $\eta_p^2$ | **Mean difference, std error** |
| aCS+/aCS- | $_{(1, 26)}$ = 7.862 | **0.009** | 0.232 | 18.433, 6.574 |
| pCS+/aCS- | $_{(1, 26)}$ = 8.614 | **0.007** | 0.249 | 17.567, 5.985 |
| pCS+/pCS- | $_{(1, 26)}$ = 6.823 | **0.013** | 0.208 | 15.578, 5.964 |
| pCS-/aCS- | $_{(1, 26)}$ = 0.221 | 0.643 | 0.008 | 1.988, 4.234 |

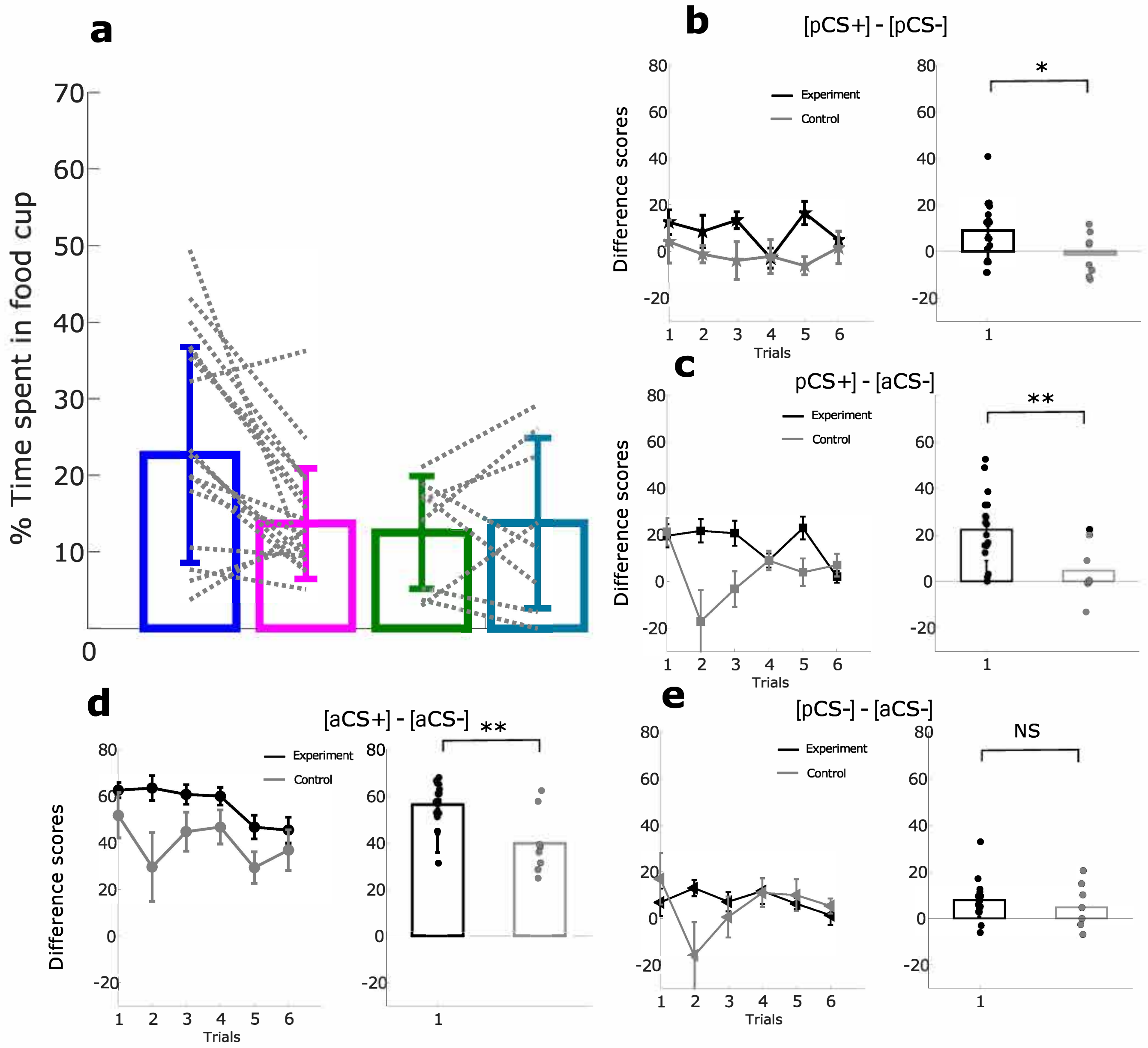


**Figure 3 - s1. Food cup occupancy in Experimental and control group.** (A) Mean percentage of Time spent in the food cup during the full 30 seconds after cue onset for experiment vs control for trial 1 to 6. (B) Difference scores of the percentage of time spent in food cup over 6 trials for the [pCS+] - [pCS-] difference scores. Bar plots show average over 6 trials of the [pCS+] - [pCS-] difference scores with dots showing the mean per rat (C) Difference of the percentage of time spent in food cup over 6 trials for the [pCS+] - [Cs-] difference scores. Bar plots show average over 6 trials of the [pCS+] - [Cs-] difference scores with dots showing the mean per rat (D) Difference of the percentage of time spent in the food cup over 6 trials for the [Cs+] - [CS-] difference scores. Bar plots show average over 6 trials of the [Cs+] - [CS-] difference scores with dots showing the mean per rat. (E) Difference of percentage of the time spent in food cup responding over 6 trials for the [pCS-] - [Cs-] difference scores. Bar plots show average over 6 trials of the [pCS-] - [Cs-] difference scores with dots showing the mean per rat. Line graph error bars indicate the SEM. (*$p < 0.05$; **$p < 0.01$; ***$p < 0.001$)

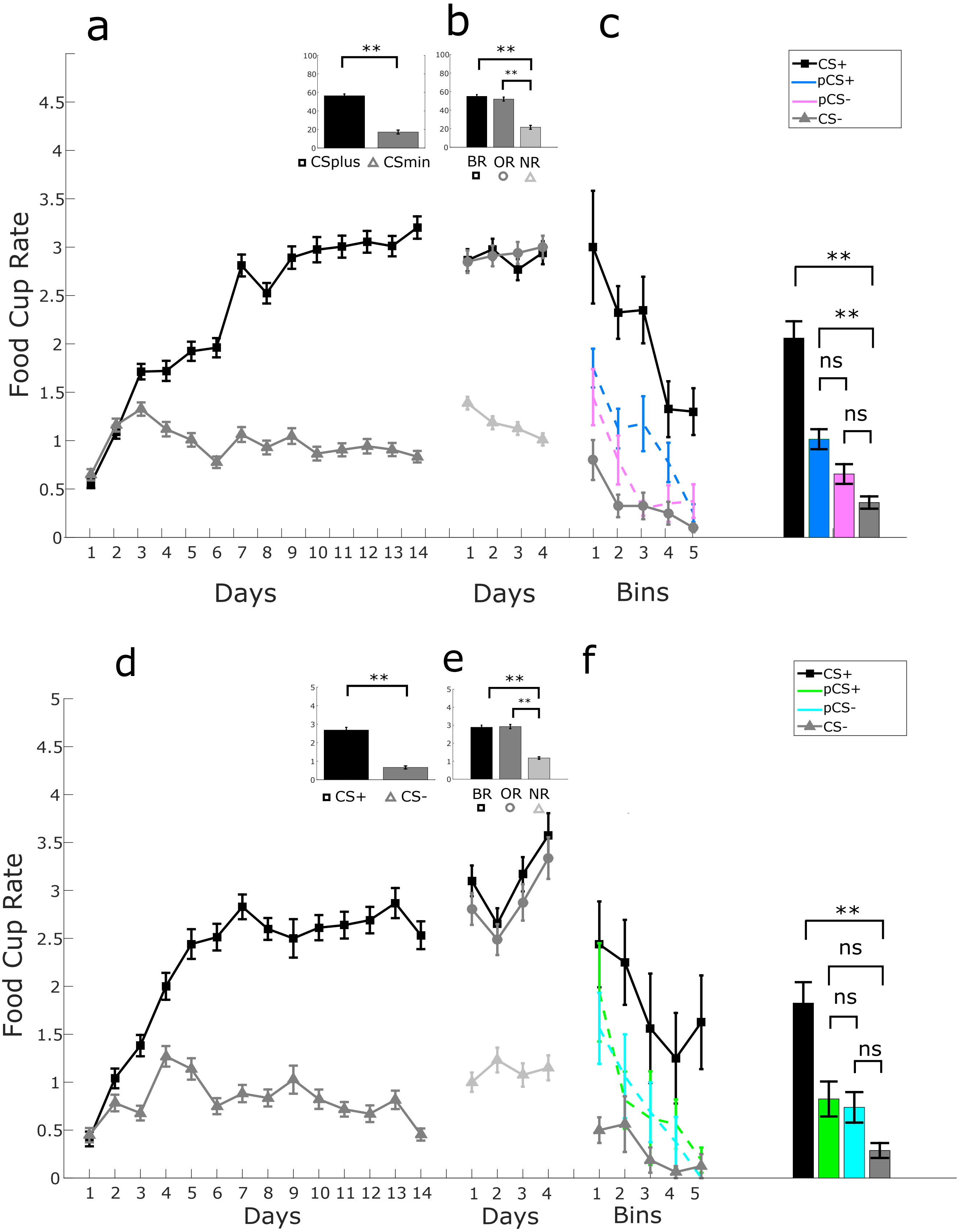


Figure S1. **Food cup rate Experimental and control group.** Experimental group. (A) Food cup rate cup for discrimination learning between aCS+ and aCS- over days. (B) Food cup rate cup for the compounds BR (aCS+, pCS+), OR (aCS+, pCS-) and NR (aCS-,pCS-) over days. (C) Percentage of time spent in the food cup during the probe trials over 10 trials. Averaged time spent in the food cup over 10 probe trials between aCS+, pCS+ (unblocked), pCS- (blocked) and aCS-. Control group. (D) Food cup rate cup for discrimination learning between aCS+ and aCS- over days. (E) Food cup rate cup for the compounds BR (aCS+,pCS+), OR (aCS+, pCS-) and NR (aCS-,pCS-) over days. (F) Percentage of time spent in the food cup during the probe trials over 10 trials. Averaged time spent in the food cup over 10 probe trials between aCS+, pCS+(unblocked), pCS- (blocked) and aCS-. Error bars indicate SEM. (*$p < 0.05$; **$p < 0.01$,***$p < 0.001$)

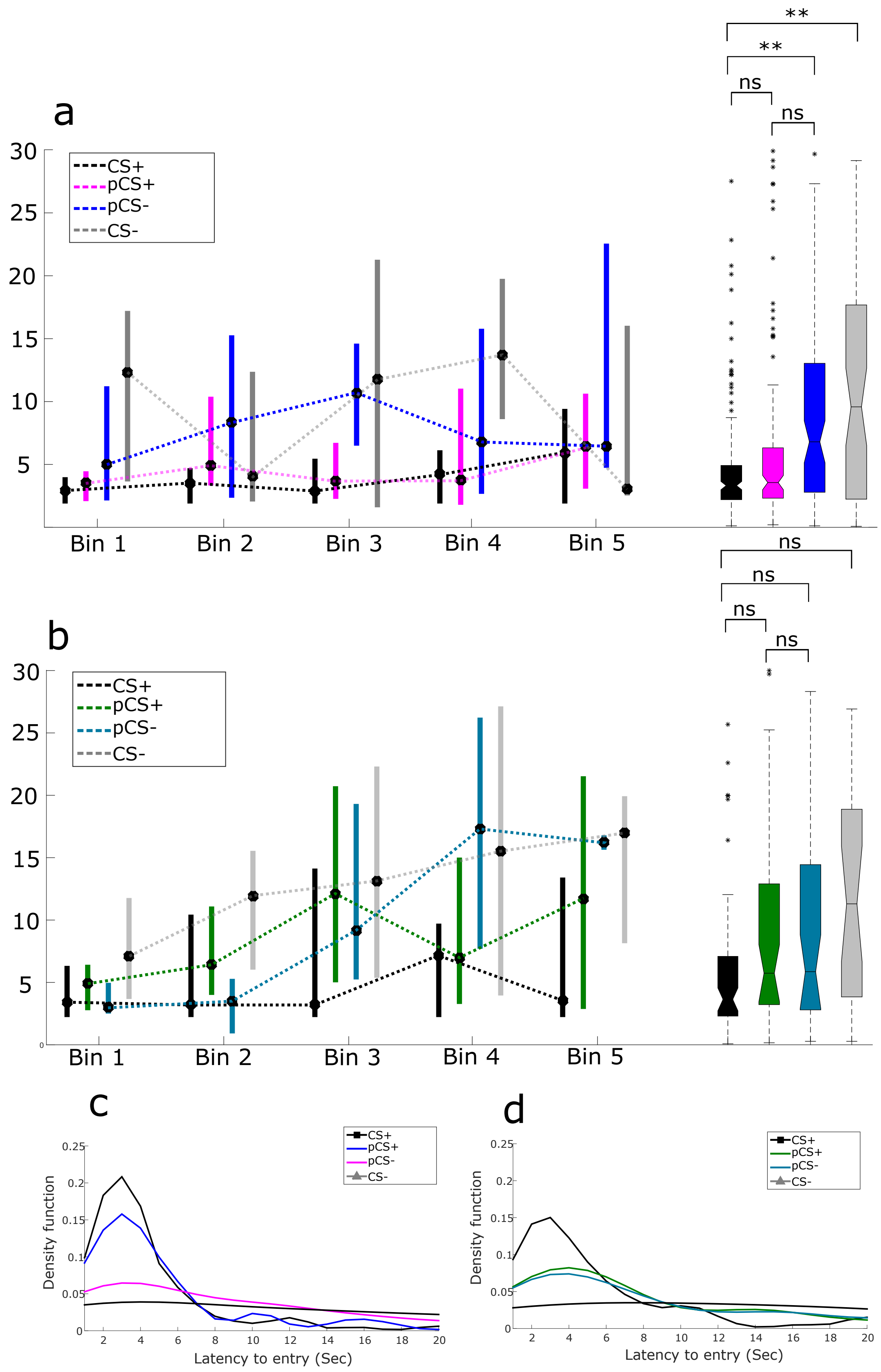


**Figure S2. Latency to entry from cue on for experimental and control group.** (A, B) Median latency to enter the food cup during the probe trials over 5 bins. Black dot indicates the median, bars display the interquartile range. Notched boxplot on the right displays the distribution per condition over all bins for experimental (A) and control (B) animals. (C, D) Kernel Density plot of the distributions of the different trial types for experimental and control animals (D). (*p < 0.05; **p <0.01)
